# NovoGlyco: mapping protein glycosylation in prokaryotes

**DOI:** 10.64898/2026.04.15.718822

**Authors:** Dinko Soic, Martin Pabst

## Abstract

Protein glycosylation in prokaryotes shows extraordinary diversity including species-specific monosaccharides, non-canonical attachment sites, and variable glycan architectures that challenge existing glycoproteomics approaches. Current strategies are largely tailored to eukaryotic systems and depend on predefined glycan databases or prior biochemical knowledge, limiting their application to microbes. Here we present NovoGlyco, a modular glycoproteomics platform for untargeted characterisation of prokaryotic protein glycosylation from shotgun proteomics data. NovoGlyco integrates de novo oxonium ion discovery, sequence tag matching, and mass offset binning to identify novel glycans, their composition, and linking chemistry. An interactive dashboard allows exploration of glycan features and modified proteins. We demonstrate the NovoGlyco platform across published glycoproteomics datasets, spanning human pathogens, Asgard archaea, and environmental enrichment cultures, and identify previously unreported flagella O-glycans in the opportunistic pathogen *Campylobacter fetus*. In summary, NovoGlyco provides a scalable framework for unbiased exploration of microbial glycoproteomes in both single-organism and metaproteomic contexts.

## INTRODUCTION

Prokaryotic protein glycosylation shows a remarkable structural diversity, only poorly captured to date^1, 2^. While mammals utilize a limited set of approximately ten different monosaccharides, prokaryotes commonly develop species-specific monosaccharide and glycan structures linked to a wider range of amino acids than the canonical serine, threonine, and asparagine residues^1, 3-5 6^. Furthermore, in bacteria protein glycosylation has been mainly studied in the context of pathogens, where glycans are virulence factors, facilitating immune evasion and infection^7^. However, the role of prokaryotic protein glycosylation extends far beyond pathogenic contexts. For example, glycans are involved in adhesion of commensal microbes to host tissues such as in the gastrointestinal system^3, 8^. Moreover, complex protein glycosylation has also been identified in strictly environmental bacteria, where glycans provide structure and support metabolic traits^2, 9^. Because prokaryotes frequently produce species-specific glycans, bacterial glycoproteins represent attractive targets for the development of novel antimicrobials and vaccines^10^.

The extensive species specific glycan diversity render traditional glycoproteomics methods unsuitable for microbial glycoproteomics studies^11^. Therefore, microbial glycoproteomics often still requires manual data curation, enrichment strategies, and carbohydrate-specific colorimetric detection^9, 12^. The chemical diversity prevents creation of glycan databases such as required for most glycoproteomic tools. Recent tools are primarily designed for mammalian glycoproteomics data. For example, GlyCombo allows identification of glycan composition from MS data using combinatorial monosaccharide analysis. However, it requires prior specification of monosaccharides^13^. GlyCounter extracts predefined oxonium, Y-type, and custom glycan fragment ions to evaluate glycan content, but relies on a predefined set of target masses rather than performing de novo identification^14^. Furthermore, glycoproteomic engines like Byonic^15^, pGlyco3^16^ and GlycanFinder^17^ are also designed for mammalian-type glycans. And while MSFragger-Glyco allows to use customized glycan databases^18^, it does not enable a completely untargeted search. Moreover, glycan database-independent search modes in Glyco-Decipher^19^, as well as GlycanFinder^17^ and pGlyco3^20^, are constrained to monosaccharides of the mammalian glycosylation pathway and conserved core structures. Furthermore, while SugarPy works without glycan databases, it still requires to define monosaccharides, the presence of unglycosylated peptide counterparts, and additional in-source fragmentation data^21^.

In contrast to database searching with exact precursor mass matches, open (or unrestricted) searching has advanced the exploration of protein modifications^22-26^. Thereby, the peptide is determined by matching fragment ions independently of the precursor ion mass. The mass difference between the matched peptide and the precursor finally provides the mass of the unspecified modification (“mass delta”)^18^. This approach, employed by Byonic wildcard search and MSFragger-Glyco, can be used for the determination of possible glycan masses, which can then be followed by focused searches, as implemented in multi-step tools such as Open-pFind^27^ and O-Pair Search^28^. While this supports the untargeted exploration of protein glycosylation in prokaryotes, bottlenecks remain that prevent large-scale studies. Firstly, the procedure often involves multi-step searches, and it includes a maximum “mass delta” that can be searched^22^. Secondly, the observed “mass delta” may not necessarily be caused by a sugar attachment but could be any type of modification. Lastly, open searches require robust peptide fragment ion coverage to enable confident sequence tag matching, which is not always given. Therefore, manual data curation and inspection are inevitable to avoid false positive and negative assignments^22^. A solution to this was recently demonstrated, where grouping of the spectra based on the presence of oxonium ions allowed subsequent extraction of glycan modification masses from precursor offsets^2^. Oxonium ions were identified by de novo annotation using an empirical sugar composition database, which allowed detection of previously unknown monosaccharides^2^. While this approach does not require any glycan reference databases, it depends on optimal fragmentation of the glycan backbone (including a detectable Y_0_ ion), and partial or too strong fragmentation may obscure the intact glycan mass.

Recently, the reliability of glyco-search engines has been also called into question due to a lack of consistent validation methods^29^. These concerns have been substantiated by a comparative study showing that even established tools report numerous spurious results for well-characterized mammalian samples^30^. If established pipelines struggle with specificity in mammalian systems with a predictable glycan structure space, the problem of spurious assignments likely magnifies for prokaryotic samples containing unexpected monosaccharides and glycan structures. Furthermore, the scarcity of glycoproteomic training data for prokaryotic samples limits deep learning-based spectral analysis tools^31, 32^. Nevertheless, a comprehensive de novo platform for the exploration of prokaryotic protein glycosylation is urgently needed to define its roles in pathogenic, commensal, and environmental microbes and their impact on human health.

Thus, we introduce NovoGlyco, a comprehensive glycoproteomics platform for identifying and characterising prokaryotic protein glycosylation directly from shotgun proteomics data. The platform includes de novo identification of oxonium ions, ultra-fast database searching, and de novo sequence tag matching for a fully untargeted glycopeptide identification, while orthogonal mass offsets provide insights into glycan composition, and linkage type. The outputs are visualised in a dashboard to support interactive exploration of identified glycoproteins. We demonstrate the platform with recently published glycoproteomics data from pathogenic bacteria, Asgard archaea and enrichment cultures from environmental microbiomes. While this enables exploration of glycoproteins in metaproteomic studies, we also identify previously undescribed flagellin O-glycosylation in the pathogen *C. fetus subsp. fetus*, which in the related C. jejuni is directly linked to host invasion and motility.

The Python code and Docker versions of OxoniumBrowser and NovoGlycoX, including documentation, are available via SourceForge: https://sourceforge.net/u/glycolab/profile/. The raw data are summarized in SI Excel Table 1 and are publicly accessible through the ProteomeXchange repositories and the 4TU.ResearchData platform (project “NovoGlyco”): https://data.4tu.nl.

## MATERIALS AND METHODS

### Outline of NovoGlyco platform

The NovoGlyco code has been implemented in Python, utilising interactive dashboard built using Plotly Dash to create an interactive browser output window that allows to investigate the results. The platform consists of two integrated modules: the first step performs identification of oxonium ions (i.e. sugar monomers) with the “OxoniumBrowser” module, and the second step performs identification of glycans and glycopeptides using the “NovoGlycoX” module. The platform was designed for high-resolution mass spectrometry data, developed and tested with data from various Orbitrap mass spectrometers (Thermo Scientific, Germany). Shotgun proteomics data acquired on other high-resolution mass spectrometers should also be compatible, provided that sufficient mass accuracy and resolution are available, and that data conversion and input parameters are adjusted accordingly. **OxoniumBrowser: de novo oxonium ion detection**. At high mass resolution and accuracy, monosaccharide-related peaks can be distinguished from amino acid-derived fragment ions^2, 33^. Shotgun proteomics data were first converted to mzML format using ThermoRawFileParser (v1.4.4) or msConvert (https://proteowizard.sourceforge.io/). Prior to oxonium ion detection, MS/MS spectra were subjected to peptide database searching using SAGE^34^ to identify and exclude unmodified peptide spectra, focusing analysis on potential glycopeptide candidates. MS/MS spectra were extracted from the mzML files using the pyteomics library (v4.7.5), retaining key spectral metadata including scan numbers, retention times, precursor m/z values, charge states, and ion intensities. Prior to analysis, extracted data underwent recalibration using known amino acid fragment peaks (147.11280, 175.11895, 201.12337, 215.13902, 228.1343, 258.1448, 292.1292 m/z) as internal calibrants. Initial global recalibration was performed using a 20-ppm mass window, followed by single-spectrum recalibration at 10 ppm tolerance. Linear recalibration parameters were determined using least squares regression, requiring a minimum of 500 matched spectra per reference peak to ensure recalibration robustness. To accommodate the extensive inter-species diversity of prokaryotic monosaccharides, two types of sugar databases are employed. The first is an empirical database curated from selected CSDB^35^ and KEGG^36^ entries and previously reported glycans^34, 37^ forming a systematic derivative space with 35 distinct monosaccharide masses. Additionally, a chemical space database consisting of over 3300 chemically plausible monosaccharide compositions within a defined elemental space C_5-14_H_4-28_N_0-2_O_2-12_S_0-1_ was constructed, enabling untargeted discovery of rare or previously unreported sugar modifications^2^. Each entry contains two diagnostic fragment masses, corresponding to the oxonium ion and its water loss mass. An equal number of random test masses scaled proportionally to the database size is included to allow empirical threshold optimization and false discovery assessment. All MS2 spectra were searched for the presence of these diagnostic ion pairs within a defined mass error tolerance of 0.001 m/z, and normalized intensity above a user-defined threshold (default 0.25% of total spectrum intensity). Detection metrics including normalized presence (percentage of spectra containing the ion pair), total spectrum counts, and both normalized and raw intensity averages across positive detections are calculated, reported and visualized through the interactive dashboard built using Plotly Dash (v2.18.1) using a summary table, a correlation plot of scans versus normalised intensity and extracted ion traces for each detected oxonium ion. The dashboard allows real-time exploration by changing detection metrics and exporting of the identified oxonium ions. **NovoGlycoX:** d**e novo glycan and glycopeptide identification**. NovoGlycoX first groups large volumes of fragmentation spectra based on the presence of (earlier identified) oxonium ions. Two complementary strategies are then applied to identify glycans and glycopeptides. The first approach generates de novo sequence tags and matches them to the reference proteome, enabling identification of the peptide backbone and mass deltas for each glycopeptide. The second approach is database-independent and calculates precursor mass offsets for all spectra (even those with poor peptide fragmentation) to infer glycan and glycan fragment masses independently of successful sequence matching. As a first step, NovoGlycoX performs a conventional database search using sagepy (v0.2.27, SAGE algorithm) to generate a focused protein database from the provided reference proteome and to identify spectra corresponding to unmodified peptides. Search parameters include static carbamidomethylation of cysteine, variable oxidation of methionine, trypsin-specific cleavage (K/R, not before P), up to 2 missed cleavages, 20 ppm precursor and fragment mass tolerances, and a minimum peptide length of 6 residues. Peptide-spectrum matches (PSMs) were filtered at 1% FDR. All unmatched MS2 spectra were then charge state deconvoluted (for charge states 2–4) and converted to singly charged m/z values. The intensity values for each fragmentation spectrum were normalised to the total ion intensity (per spectrum), and spectra containing oxonium ion/water loss pairs above a user-defined intensity threshold (default 0.1% of total intensity) were flagged. Next, these spectra were subjected to de novo sequencing using DirecTag^*38*^ to generate sequence tags (default length 5 amino acids). The top ten tags, ranked by their scoring metrics, were matched against in silico-generated peptides from the focused reference proteome using a dictionary-based lookup approach, with the option to restrict matches to peptides containing serine, threonine, or asparagine as potential glycosylation sites. Valid matches required complete tag presence within the candidate peptide sequence and the calculated peptide mass to be at least 100 Da smaller than the precursor mass. Potential glycopeptide matches are optionally validated by the presence of unmodified peptide ions (Y0) within 20 ppm mass tolerance. For peptides >2000 Da where the singly charged peptide is not found, the code checks for the presence of the doubly charged variant. For each spectrum, the peptide-protein combination with the largest number of sequence tag matches was reported as the identification, where confidence increases with the number of tag matches. Reported results include the identified peptide sequence, the corresponding mass delta, and the oxonium ions detected in the spectrum. For visualization, mass deltas associated with each oxonium ion are binned into histograms and displayed as bar graphs. True glycan modifications were revealed as recurring mass deltas across different peptide sequences. Furthermore, the second approach determines precursor mass offsets by calculating for each oxonium ion-containing spectrum the mass difference between the precursor and all fragment ions above a defined mass threshold (default, 500 or 750 Da). The preferred fragmentation of the glycan chain gives rise to a series of glycopeptide Y ions (sole glycan fragments)^*2, 39-41*^ which aids in reconstructing the oligosaccharide structures^2, 16^. Subtracting these fragment ion masses from the precursor (precursor offsets) reveals both the intact glycan mass and glycan-derived fragments, offering a database-independent method for identifying glycan compositions^2^. Complementary peptide offsets are calculated from the mass differences between the observed unmodified peptide ion (Y0) and higher mass fragments, representing sequential monosaccharide additions to the unmodified peptide. This dual analysis provides complementary evidence for glycan structure, with precursor mass offsets reflecting sequential monosaccharide losses from the intact glycopeptide, and peptide mass offsets revealing stepwise glycan assembly, including the linking sugar. For visualization, precursor and peptide offsets for each oxonium ion are binned into histograms and displayed as bar graphs. The bar graphs and histograms are displayed in an interactive dashboard built using Plotly Dash (v2.18.1) enabling interactive exploration and interpretation of the data. Additionally, a heatmap displays oxonium ion correlation patterns across spectra, where the colour represents the degree of correlation between different oxonium ion pairs. For glycan composition analysis visualization, the dashboard provides three interlinked histogram types: mass deltas (differences between precursor and peptide masses), precursor offsets (mass losses from intact glycopeptides), and peptide offsets (mass additions to peptide backbone). All mass frequencies were analysed with a bin width of e.g. 1 or 0.1 Da, effectively clustering similar mass values while maintaining sufficient resolution to distinguish different monosaccharide combinations. User selection of distinct bins in any histogram triggered automatic updates in related plots, showing corresponding distributions in the other metrics. For selected glycan mass modifications, the dashboard also featured interactive glycoprotein candidate table displaying supporting evidence including individual peptide-spectrum matches with scoring metrics and oxonium ion annotations. All visualizations support real-time filtering and selection, and results can be exported in both detailed tabular format and PeptideShaker^*42*^ compatible formats for further analysis and integration with standard proteomics workflows. NovoGlyco is an open-source Python platform that is freely available under the Apache License 2.0, and the code and Docker version for both modules OxoniumBrowser and NovoGlycoX including all necessary dependencies and tools are available via SourceForge, including documentation https://sourceforge.net/projects/novoglycox/files/ and https://sourceforge.net/projects/oxoniumbrowserx/files/. **Microbial proteomics datasets**. A summary of the analyzed species, corresponding raw file names, reference databases, and publicly available datasets is provided in the SI Excel Table S1. Shotgun proteomics raw data for Campylobacter jejuni DSM 27585 and Escherichia coli BL21 are available via the ProteomeXchange project PXD076598, together with the reference proteomes UP000000799 and UP000000625, respectively. Proteomic data for Prometheoarchaeum syntrophicum and Sulfolobus acidocaldarius were obtained from ProteomeXchange projects PXD051006 and PXD051007^23^, and the corresponding reference proteomes were retrieved from UniProtKB (UP000321408 and UP000001018_330779, respectively). Proteomics datasets for Acinetobacter baumannii (PXD018587)^22^, Campylobacter fetus subsp. fetus (PXD018587)^22^, Porphyromonas gingivalis (PXD028120)^24^, Tannerella forsythia (PXD026989)^43^, and Ns. viennensis (PXD056844)^44^ were processed using UniProtKB reference proteomes UP000197394, UP000000760, UP000008842, UP000005436, and UP001059771, respectively. The Haloferax volcanii proteome (PXD021600^2^) was analysed using the UniProtKB reference database UP000008243_30980. From the same project (PXD021600^2^), proteomics data for “Candidatus Kuenenia stuttgartiensis” and “Candidatus Brocadia sapporoensis” were analysed using custom reference databases. Control datasets for Escherichia coli K12 and Saccharomyces cerevisiae were also obtained from PXD021600^2^ and analysed using UniProtKB reference proteomes UP000000625_83333 and UP000002311_559292, respectively. Detailed experimental protocols are provided in the corresponding PRIDE entries and referenced publications. The collection of reference data is summarized in SI Excel Table 1 and is available via the 4TU.ResearchData repository: https://data.4tu.nl (project “NovoGlyco”).

## RESULTS

The NovoGlyco platform enables fully untargeted discovery of prokaryotic protein glycosylation from shotgun proteomics data, and it consists of two modules called “OxoniumBrowser” and “NovoGlycoX (explore)” (Figure 1). The OxoniumBrowser identifies sugar-derived fragment ions (oxonium ions), whereas NovoGlycoX determines the intact glycan mass, its composition, linking-sugar/type, and corresponding glycoproteins. To account for the unpredictable diversity of monosaccharides in prokaryotes, OxoniumBrowser compares all low-mass fragment ions against either an empirical sugar database—comprising a systematic range of representative pentose, hexose, heptose, and ulosonic acid derivatives together with diagnostic monosaccharide marker fragments—or a chemical composition space containing over 3300 chemically plausible compositions, which enables the untargeted discovery of rare sugars^2^. To ensure correct annotation of oxonium ions, it was expected to also observe water-loss fragment(s), because sugar ions show sequential dehydration during fragmentation. Simultaneously, searches are performed against an equal set of random oxonium masses to estimate the degree of random annotations. Finally, identified oxonium ions are clustered by their co-occurrence across fragmentation spectra, indicating which monosaccharides belong to the same glycan structure. Next, in NovoGlycoX, the identified oxonium ions guide the identification of the intact glycan and the associated glycoproteins. First, spectra containing oxonium ions are de novo sequenced to generate sequence tags, that are matched against a reference sequence database to identify the corresponding tryptic peptide. The mass delta between the matched tryptic peptide and the observed precursor is then reported as glycan modification. Binning these mass deltas into histograms generates the so called “mass-delta spectra” with distinct peaks indicating glycan masses. Mass deltas can be grouped according to occurring oxonium ions, which gives for each oxonium ion cluster a separate mass-delta spectrum, adding compositional information to each glycan mass peak. Finally, NovoGlycoX provides an interactive visualisation of the glycan and fragment mass histograms, the associated glycopeptides and glycoproteins. However, sequence-tag matching requires a good fragment ion coverage which is not always achieved for microbial samples. Therefore, NovoGlycoX additionally applies a parent mass offset approach as an orthogonal strategy to identify glycan masses and infer glycan sequences, which has recently been demonstrated for both small mass modifications and prokaryotic glycopeptides^2, 45^. For every oxonium ion containing fragmentation spectrum, the precursor mass is subtracted from each fragment ion. Binning the resulting parent offsets into histograms generates mass-delta spectra, in which recurring mass differences, such as those arising from sequential monosaccharide losses, appear as frequency peaks. These reveal the intact glycans and their fragments, therefore also providing insight into glycan sequence, the linking sugar, and linkage type. This alternative approach enables determination of glycan masses without reference sequence databases or de novo sequence tags. It has been shown effective for prokaryotic glycosylation, where glycoform diversity within a given strain is typically limited^2^. Finally, both tools allow interactive exploration of the identified oxonium ions, glycans, glycopeptides, and glycoproteins through a browser-based user interface (SI Figures 1 and 2).

**Figure 1.**
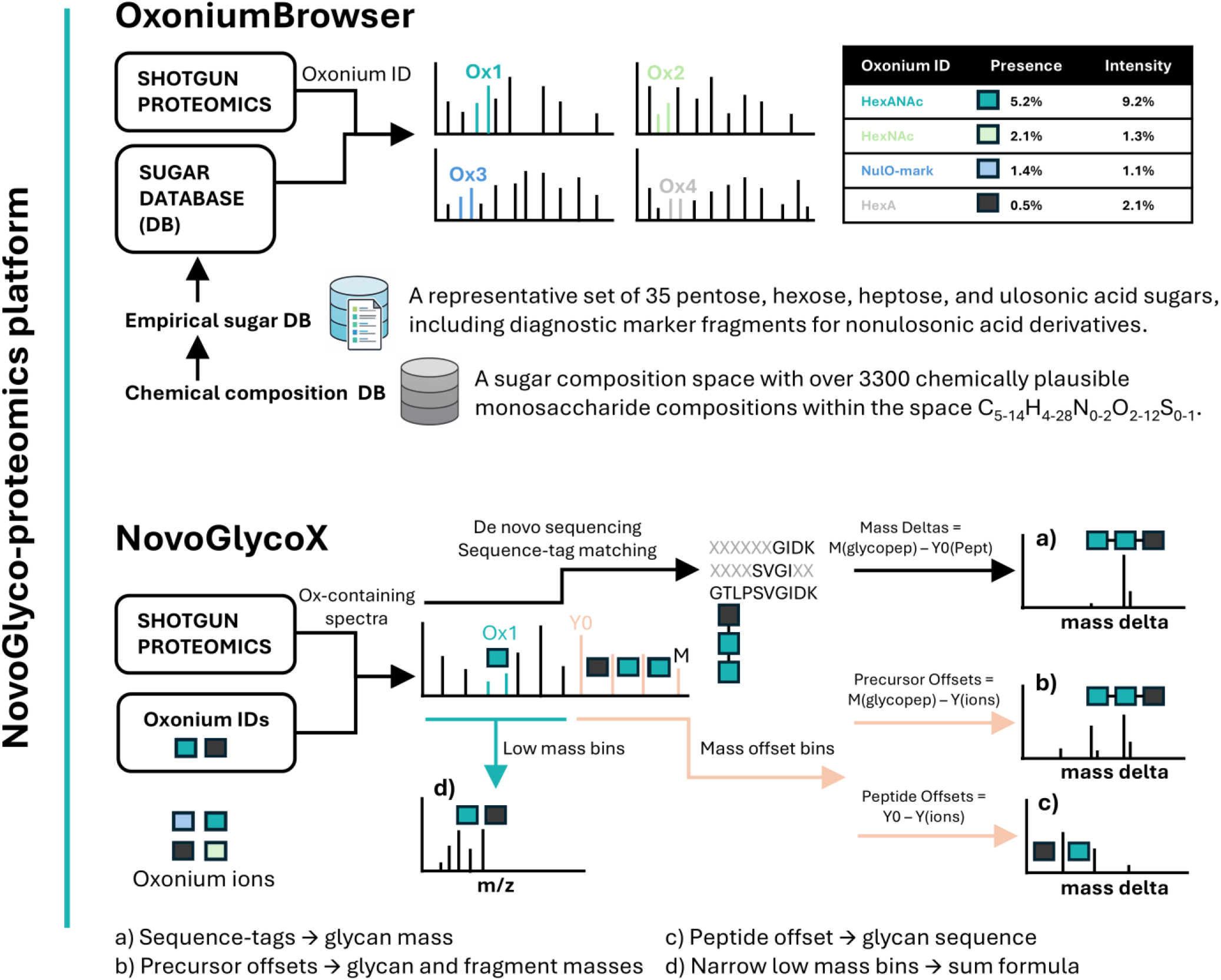
Schematic overview of the NovoGlyco microbial glycoproteomics platform. The platform consists of two modules, OxoniumBrowser and NovoGlycoX, which together enable fully untargeted exploration of protein glycosylation in microbes from shotgun proteomics data. Both modules import shotgun proteomics data in mzML format along with a reference proteome. The OxoniumBrowser additionally uses either an empirical monosaccharide database or a large chemical composition space (>3,300 candidate sugar compositions). Results from both modules are displayed in interactive dashboards (shown in SI DOC Figures 1 and 2). OxoniumBrowser reports identified oxonium/water loss pairs in tables, correlation plots, and extracted ion chromatograms. NovoGlyco displays histograms and tables for observed glycan masses, sequence informing fragments, linkage types, and modified proteins. **OxoniumBrowser workflow** Low-mass fragment ions, acquired at high mass accuracy and resolution, are matched against a sugar database to identify oxonium/water loss pairs. An equal number of random oxonium masses are included to estimate false matches. Identified oxonium ions are then clustered by co-occurrence to group monosaccharides belonging to the same glycan. **NovoGlycoX workflow** (a) Spectra containing oxonium ions are subjected to de novo sequencing and resulting peptide sequence tags are matched to a reference proteome. Mass differences between peptides and precursors are reported as mass deltas, which are binned per oxonium ion to generate pseudo-mass delta spectra representing intact glycan masses. (b) Parent mass offsets, calculated by subtracting fragment ions from the precursor mass, provide intact glycan masses along with additional glycan fragments. Diagnostic peaks around the intact glycan provide information on linkage type (N-versus O-linked). (c) Peptide mass offsets, calculated by subtracting fragment ions from the peptide (Y_0_ ion), produce glycan sequence spectra, starting from the linking sugar attached to the protein backbone. (d) Finally, summing spectra containing a given oxonium ion at narrow bin size enables determination of accurate mass and sum formula for previously unidentified fragment ions.

### NovoGlyco-proteomics across a broad taxonomic group of microbes

We demonstrate the NovoGlyco platform using shotgun proteomics datasets from public repositories spanning a broad range of taxonomic groups. These include the pathogenic bacteria Campylobacter jejuni, Campylobacter fetus subsp. fetus, Porphyromonas gingivalis, Tannerella forsythia, and Acinetobacter baumannii, the archaea Prometheoarchaeum syntrophicum, Sulfolobus acidocaldarius, and Haloferax volcanii, as well as environmental *Ca*. Kuenenia stuttgartiensis and *Ca*. Brocadia sapporoensis enrichment cultures. The mass spectrometric raw files, reference databases, and ProteomeXchange project identifiers are provided in the SI Excel Table S1. **OxoniumBrowser: de novo oxonium ion discovery**. First, we used OxoniumBrowser to de novo identify oxonium ions in the fragmentation spectra of the 13 species (Figure 2). These included non-glycosylated E. coli K12 and glycosylated yeast (S. cerevisiae) as controls, to optimize parameter settings, such as mass error tolerance, detection thresholds, and post-processing filters. Overall, the identified oxonium ions were largely species-specific, as shown in Figure 2 and summarized in SI Tables S4 and S5. As expected, in E. coli K12, no significant oxonium ions were detected apart from low-level hexose fragments, which are further discussed in the following section^2^. In contrast, S. cerevisiae showed strong and highly co-occurring HexNAc and hexose oxonium ions, consistent with the canonical N- and O-linked high-mannose glycans in yeast^46^. However, oxonium ions from the other microbial species were highly diverse, including many unconventional derivatives with multiple amine groups and a wide range of nitrogen and oxygen modifications beyond common methylation and acetylation. Also, oxonium ions derived from amino-modified monosaccharides generally showed much higher abundance than those from hydrophobic or acidic monosaccharides as described recently^1, 2^. Although this favours the detection of amino-modified sugars, most nitrogen-free oxonium ions were detected when lowering detection thresholds in the OxoniumBrowser (i.e. <0.1% of max MS2 peak intensity).

**Figure 2.**
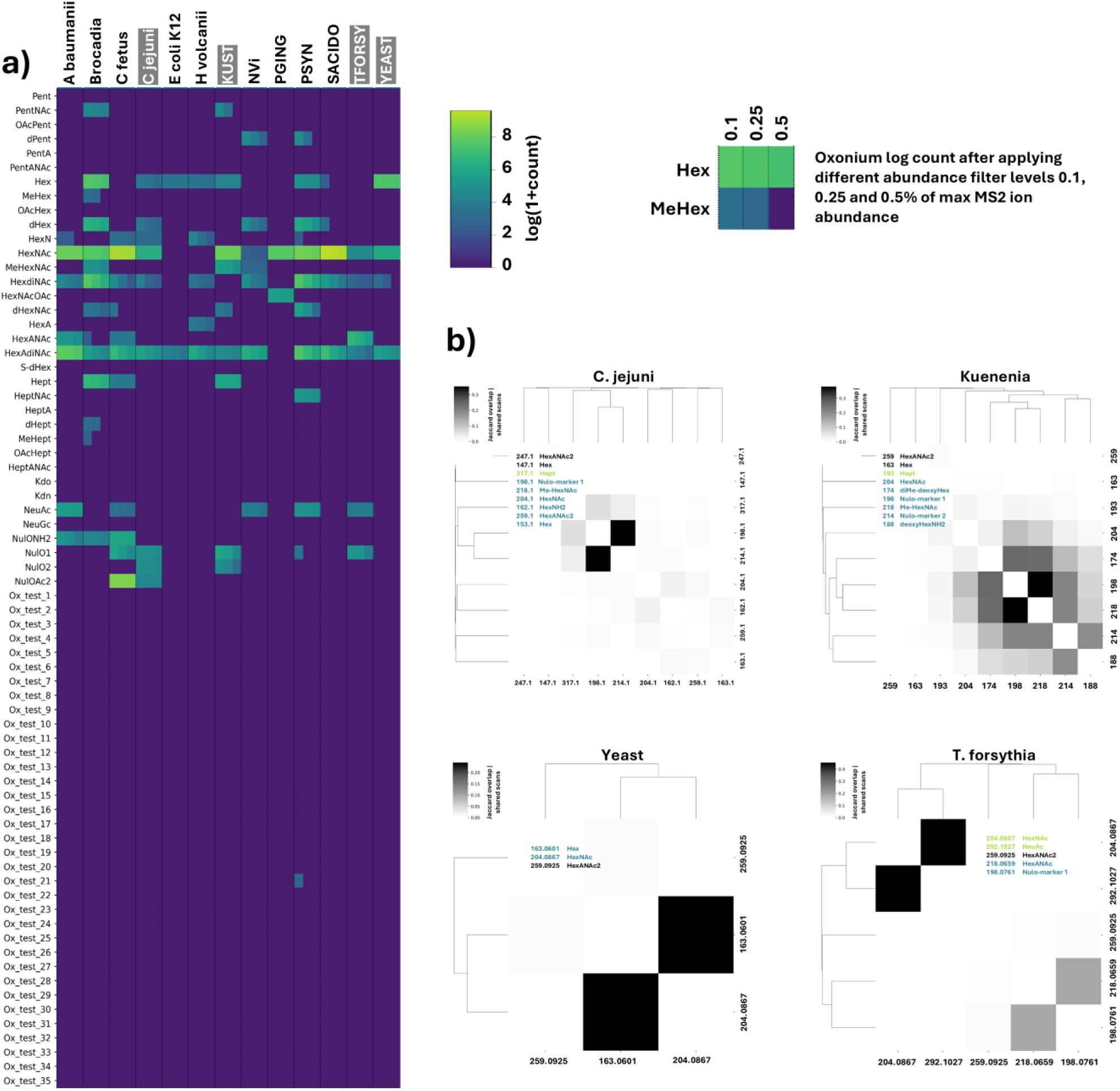
Species-specific oxonium ion profiles observed in shotgun proteomics datasets as identified by OxoniumBrowser. a) Detected oxonium ions across the empirical monosaccharide database space (including equal number of randomly generated oxonium ions for validation). The heatmap displays log(1+count) values for each oxonium ion at three detection thresholds (0.1%, 0.25%, and 1% of the maximum MS/MS peak abundance). Samples from left to right: Acinetobacter baumannii (A. baumannii), Ca. Brocadia sapporoensis enrichment (Brocadia), Campylobacter fetus subsp. fetus (C. fetus), Campylobacter jejuni (C. jejuni), Escherichia coli K12 (E. coli K12), Haloferax volcanii (H. volcanii), Ca. Kuenenia stuttgartiensis (KUST), Nitrososphaera viennensis (NVi), Porphyromonas gingivalis (PGING), Prometheoarchaeum syntrophicum (PSYN), Sulfolobus acidocaldarius (SACIDO), Tannerella forsythia (TFORSY), and Saccharomyces cerevisiae (YEAST). A summary of the identified oxonium ions, analyzed mass spectrometric raw files, and corresponding ProteomeXchange identifiers is provided in SI Excel Table 1. b) Clustering of identified oxonium ions based on Jaccard overlap (shared scan counts) for Saccharomyces cerevisiae (Yeast), Tannerella forsythia, Campylobacter jejuni, and (Ca) Kuenenia stuttgartiensis. Colours and coloured boxes indicate oxonium ions that cluster together and are inferred to originate from the same glycan structure.

Interestingly, HexNAc oxonium ions were detected in 12 of the 13 organisms, albeit five species including H. volcanii, T. forsythia, Ca. Kuenenia stuttgartiensis, and Nitrososphaera viennensis produce glycans that lack HexNAc residues. However, in T. forsythia and A. baumannii, the HexNAc ions clustered separately from the known microbial glycan monosaccharides and instead grouped with NeuAc fragments. These strains were cultured on media containing animal proteins (e.g., calf serum), suggesting that the detected HexNAc/NeuAc derive from residual medium glycoproteins rather than microbial glycans. Furthermore, Ca. Kuenenia stuttgartiensis produces two distinct glycan structures, and only one contains HexNAc residues, while the other consists of repeating heptose units. This is also reflected in the cluster graph, where the heptose-derived oxonium ions (m/z 193.0707) group separately from the HexNAc-core glycan. Several of the investigated species are known to decorate their glycans with nonulosonic acids (NulOs). However, NulO derivatives are challenging to detect because they are typically modified and difficult to recognize by mass alone. However, NulOs generate characteristic secondary fragments, such as the NulO1 (m/z 198.0761) and NulO2 (m/z 214.1074) marker ions^2^. These were indeed consistently observed whenever NulO modifications were expected, for example in C. jejuni, where NulOs modify flagellin proteins^47-49^. Accordingly, C. jejuni showed two distinct oxonium ion clusters: one corresponding to the conserved HexNAc-containing N-glycan and another to flagellin-associated NulO modifications^48-50^. Similar signals were observed in C. fetus, suggesting a comparable, yet undescribed NulO modification in this organism. Notably, some nitrogen-rich sugars remain difficult to distinguish from peptide- or amino acid-derived fragment ions even at very high mass resolution. For example, BacdiNAc (m/z 211.1077, not shown) and HexAdiNAc (m/z 259.0925) overlap substantially with peptide related fragments and are therefore unsuitable as standalone oxonium ion markers. However, in *C. jejuni* and *C. fetus*, the presence of BacdiNAc was accompanied by additional sugar fragments (e.g., hexosamine, m/z 162.0761), which could serve as proxies for its detection. In addition, supposedly rare monosaccharides, such as heptose (m/z 193.0707) and HexNAcA (m/z 218.0659) were detected in multiple organisms, including Brocadia and Kuenenia enrichments, C. fetus, A. baumannii, and T. forsythia. Furthermore, N. viennensis^44^ and T. forsythia^51^ produce exceptionally diverse amino-modified and oxidized glycans, devoid of the common HexNAc residues and would be missed by conventional approaches, without enrichment or multi-stage analysis.

### NovoGlycoX: oxonium-directed discovery of glycans, composition, linkage type, and glycoproteins

The identified oxonium ions subsequently guide identification of the glycans. For every organism, spectra containing the identified oxonium ions were subjected to de novo sequencing, sequence tag matching and calculation of parent mass offsets, where resulting mass deltas were binned into mass-delta spectra. The interactive output allows furthermore exploration of the intact glycan mass, glycan sequence, sugar linkage type and glycoproteins. Figure 3 (panel a) demonstrates the advantage of the oxonium-guided sequence tag and mass offset approaches over the conventional (unguided) open search approaches. The open search simply provides mass deltas, and without additional experiments, it is difficult to determine which of them are glycan-related. Pre-grouping fragmentation spectra based on the presence of oxonium ions enables sequence-tag matching and generation of mass-offset spectra, in which the observed mass deltas correspond to glycans containing these oxonium ions. The interactive NovoGlycoX dashboard further allows investigation of fragment ions associated with these glycan peaks, revealing their compositional sequence and linkage type. Additionally, restricting the mass tag search to spectra that contain the Y0 ion provides further confidence in the mass deltas. In *Ns. viennensis*, for example, focusing on fragmentation spectra that include HexdiNAc peaks (m/z 246.12) reveals a glycan with 1650 Da, composed of highly oxidized amino-sugars^44^. *Ns. viennensis* is a terrestrial ammonia-oxidizer where the nitrogen rich compounds (such as their amino-sugars) might have evolved to enhance nitrogen storage and reuse as a survival strategy^44^. The additional peaks in the parent offset spectrum and a zoom onto the satellite peaks around 1650.65 Da further reveal the glycan sequence and linkage type (N- or O-glycan). As reported previously for mammalian type glycans^39, 52^, HexNAc-containing N-glycans show a characteristic HCD fragmentation pattern with +17 and −84 Da shifts, reflecting ammonia loss from asparagine and cross-ring cleavage of the linking sugar. Similarly, the N-linked glucose/glucuronic acid glycan in *H. volcanii* shows a characteristic +17 and −42 Da fragmentation pattern, derived from the glucose linking sugar, with the −42 shift reflecting the absence of N-acetylation^2^. This pattern was consistently observed across the microbial datasets analyzed in this study. Panel b) of Figure 3 furthermore illustrates how glycan composition and sequence can be resolved from mass offset data, exemplified with the glycoproteomic data for *P. syntrophicum*^*23*^. This organism is an Asgard archaeon of particular interest due to its close phylogenetic relationship to eukaryotes^53, 54^. When processing the spectra containing the major oxonium ion (HexNAc), NovoGlycoX recovered the same three glycoforms as reported previously by Nakagawa and co-workers^*23*^, with masses of 801, 909, and 1213 Da. These masses were also detected in the parent mass offset spectra, although glycan fragmentation provides additional (smaller mass delta) peaks. To determine the composition and sequence of these three glycans, the interactive dashboard was used to generate the parent and peptide offset mass deltas of spectra representing each glycan separately. Interestingly, this showed a common linking sugar (i.e. 304.13 Da, likely a serine- or threonine-modified HexNAc derivative^*23*^) extended by a Pent-Hex moiety, with differing terminal extensions. For example, the 801 Da glycan contained an additional 303.10 Da, the 909 Da glycan an additional 311.08 Da, and the 1213 Da glycan contained both of these terminal sugars. A zoom onto the glycan 909 shows only an additional +18 satellite peak, consistent with the O-linkage of these glycans. Notably, despite its evolutionary proximity, the N-glycan core structure of P. syntrophicum shows little similarity to canonical eukaryotic N-glycans. In contrast, the more distantly related crenarchaeon Sulfolobus acidocaldarius produces glycans that more closely resemble those of eukaryotes as they incorporate a chitobiose core disaccharide^55, 56^. Panel c) in Figure 3 shows the detection of NulO type O-glycans on flagellin proteins in *C. jejuni* and *C. fetus fetus*, the latter not described previously. For C. jejuni, the peptide mass offsets reveal mono- and dimers of NulO derivatives, including NulOAcAm, NulOAc_2_, and NulOAcMeGA, as well as low-abundance variants also described recently by van Ede et al (van Ede et al., bioRxiv 2026). These NulOs comprise mixtures of Pse and Leg stereoisomers and are exclusively O-linked to FlaA and FlaB, consistent with the characteristic +18 Da satellite peak^49, 57^. Interestingly, OxoniumBrowser indicated similar NulOAc_2_ modifications in *C. fetus fetus*. However, the mass-offset spectra show more prominent dimer peaks, whereas sequence-tag matching predominantly identified NulOAc_2_ trimers modifying the flagellin proteins A0RRD1 and A0RRD0 (annotated spectra are shown in SI Doc Figures 5–9). Although not previously reported in C. fetus, flagellin NulO modifications have been identified in related pathogens, including C. jejuni and H. pylori, where they serve as important virulence factors and are required for proper flagellar assembly and motility^7, 48, 49, 58^. This suggests that the identified flagellin glycans in *C. fetus* may also represent important virulence factors, with potential implications for host interactions. However, in the reanalyzed dataset, the identified NulO-modified peptides eluted at the very end of the chromatographic gradient. Therefore, additional NulO oligomers or other variants may be present but were not captured under these experimental conditions.

**Figure 3.**
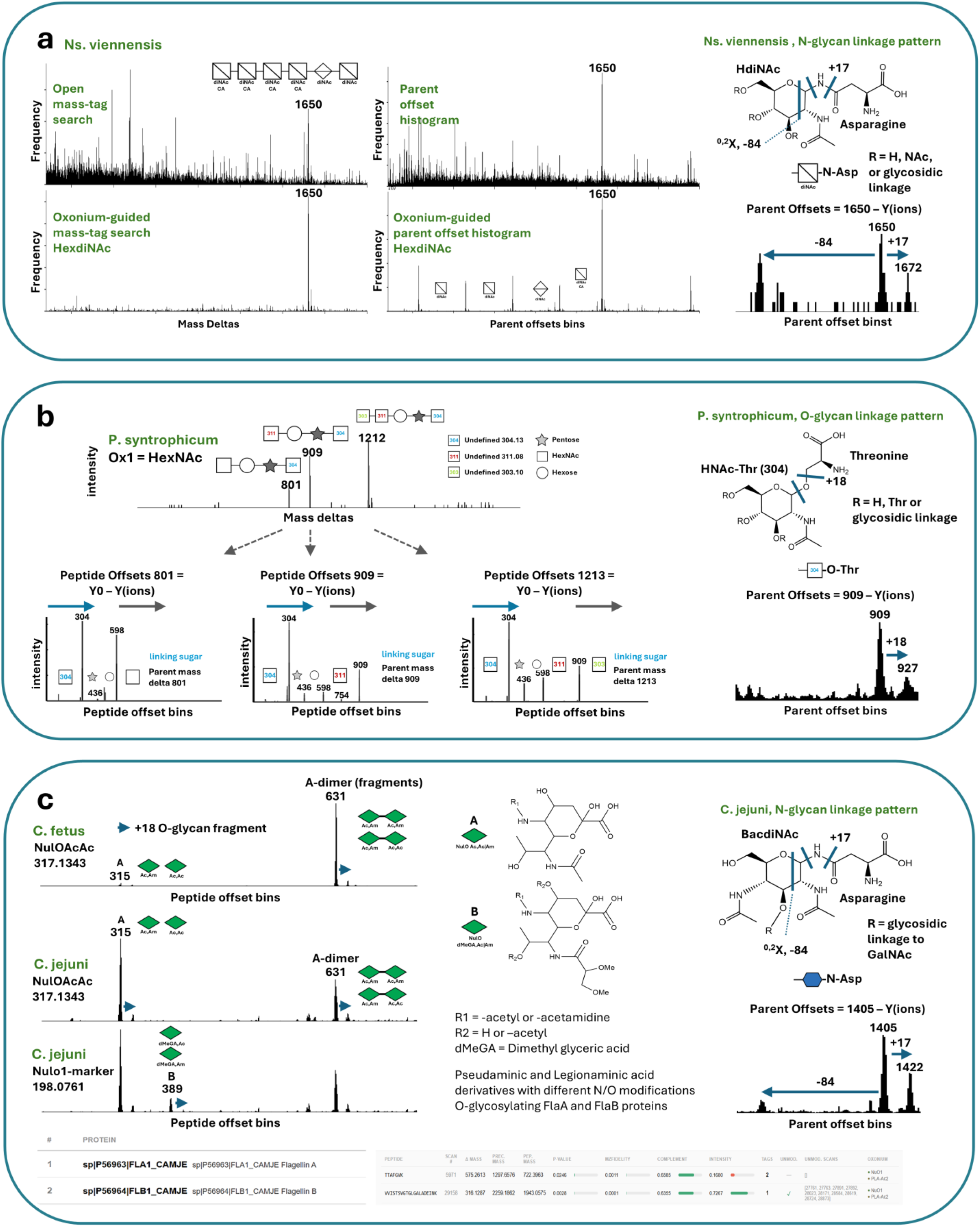
NovoGlycoX oxonium-focused search, glycan sequence and linkage-type determination. Panel a) Comparison of oxonium-guided analysis and open search for N. viennensis. Unguided open searches yield numerous unspecific mass deltas, whereas oxonium-guided grouping (e.g., HexdiNAc, m/z 246.12) produces clean sequence tag and mass offset spectra, directly revealing a single glycan of 1650 Da. Satellite peaks around the glycan mass show characteristic +17/−84 Da signature fragments, consistent with N-linked HexNAc-type glycans. NovoGlycoX was run with an oxonium ion intensity threshold of 0.1%, using a bin size of 0.1 Da (linkage type figure 1Da), a minimum offset mass of 750, and peptide mass validation disabled. Panel b) Together, mass offset and peptide offset spectra enable reconstruction of glycan architecture as exemplified for P. syntrophicum. Three glycoforms (801, 909, and 1213 Da) are identified which can be selected for further exploration in the interactive output window. Investigating the peptide offset histograms show that these glycans share a common 304.13 Da linking sugar extended by a Pent–Hex core. Differences arise from distinct terminal extensions, either +303.10 Da, +311.08 Da, or both. The +18 Da satellite peak around the glycan masses confirms their O-linkage. NovoGlycoX was run with an oxonium ion intensity threshold of 0.01%, using a bin size of 0.01 Da (linkage type graph 1Da), a minimum offset mass of 500, and peptide mass validation disabled. Panel c) Identification of NulO-type O-glycans on flagellins in C. jejuni and C. fetus fetus. In C. jejuni, peptide mass offsets show mono- and dimeric NulO derivatives (e.g., NulOAcAm, NulOAc_2_), consistent with known Pse/Leg-modified Fla proteins. In C. fetus fetus, similar patterns are observed, with dominant NulOAc_2_ trimers modifying flagellins (A0RRD1, A0RRD0), representing previously undescribed glycosylation. These findings suggest conserved flagellin glycosylation linked to motility and virulence, although additional higher-order NulO oligomers may be present. NovoGlycoX was run for CJ with an oxonium ion intensity threshold of 0.25%, and for Cff with a threshold of 0.1%, using a bin size of 0.1 Da (linkage type figure 1Da) and minimum offset mass of 500, and peptide mass validation disabled.

Figure 4 provides an overview of the dozens of glycans and glycoforms that were identified across the analyzed microbes with the NovoGlycoX module. The left panels show spectra from sequence tag matching, while the right panels display spectra derived from parent mass offsets. When OxoniumBrowser detected multiple glycan clusters within the same organism, mass deltas from one representative oxonium ion per cluster were plotted in a single spectrum. Both approaches (sequence-tag matching and parent mass offsets) generally identify the same glycans, but with notable differences. Firstly, parent mass offsets provide additional fragment masses from the reducing end, enabling reconstruction of glycan composition and sequence. Secondly, when peptide backbone fragmentation is poor, sequence-tag matching may yield missing or low-frequency mass deltas. For example, the H. volcanii and the Campylobacter glycans were barely detected with the sequence tag matching approach, whereas the parent offset approach provided clear glycan peaks. Optimizing collision energies to balance peptide backbone and glycan fragmentation can enhance detection with either approach (SI DOC Figure 3). Thus, while de novo sequence tag matching identifies the intact glycans and modified peptides, parent offset mass delta provide complementary information on glycan composition and the linking sugar type. Overall, for all microbes, the observed glycans were consistent with published data, with several unique features worth noting. For example, in H. volcanii the predominant glycoform was the recently detected tetrasaccharide (Asn-Hex-HexA-HexA-MeHexA, 704 Da), whereas the larger pentasaccharide (866 Da) was observed only at very low abundance^2, 59-61^. Furthermore, only a single type of glycan was detected, consistent with the high-salinity growth conditions (>3.5 M NaCl)^61, 62^. For Campylobacter fetus, previously unreported O-linked NulO flagellin modifications were detected (as described above and shown in Figure 3c), similar to those in Campylobacter jejuni. In addition, the known N-linked glycoforms at 1243.51 Da and 1202.48 Da were observed, together with formylated (+28 Da) and truncated variants. However, it is not clear whether the formylation represents a native modification or resulted from formic acid exposure during sample preparation or storage^63^. Finally, analysis of Sulfolobus acidocaldarius confirmed the previously characterized branched and sulfated N-glycan, with the predominant 1118.35 Da glycoform and a truncated 956.25 Da variant. In Acinetobacter baumannii, the characteristic 1030.35 Da pentasaccharide and its methylated and acetylated derivatives were also detected as described earlier^64^. For Porphyromonas gingivalis, the three major known O-linked glycoforms, 1352.45, 1394.45 (+1 Ac), and 1436.45 Da (+2 Ac), were observed^24^. Similarly, in T. forsythia, NovoGlycoX correctly resolved the complex O-glycan composed of a glucuronic acid/galactose core with a terminal NulO modification^51, 65^.

**Figure 4.**
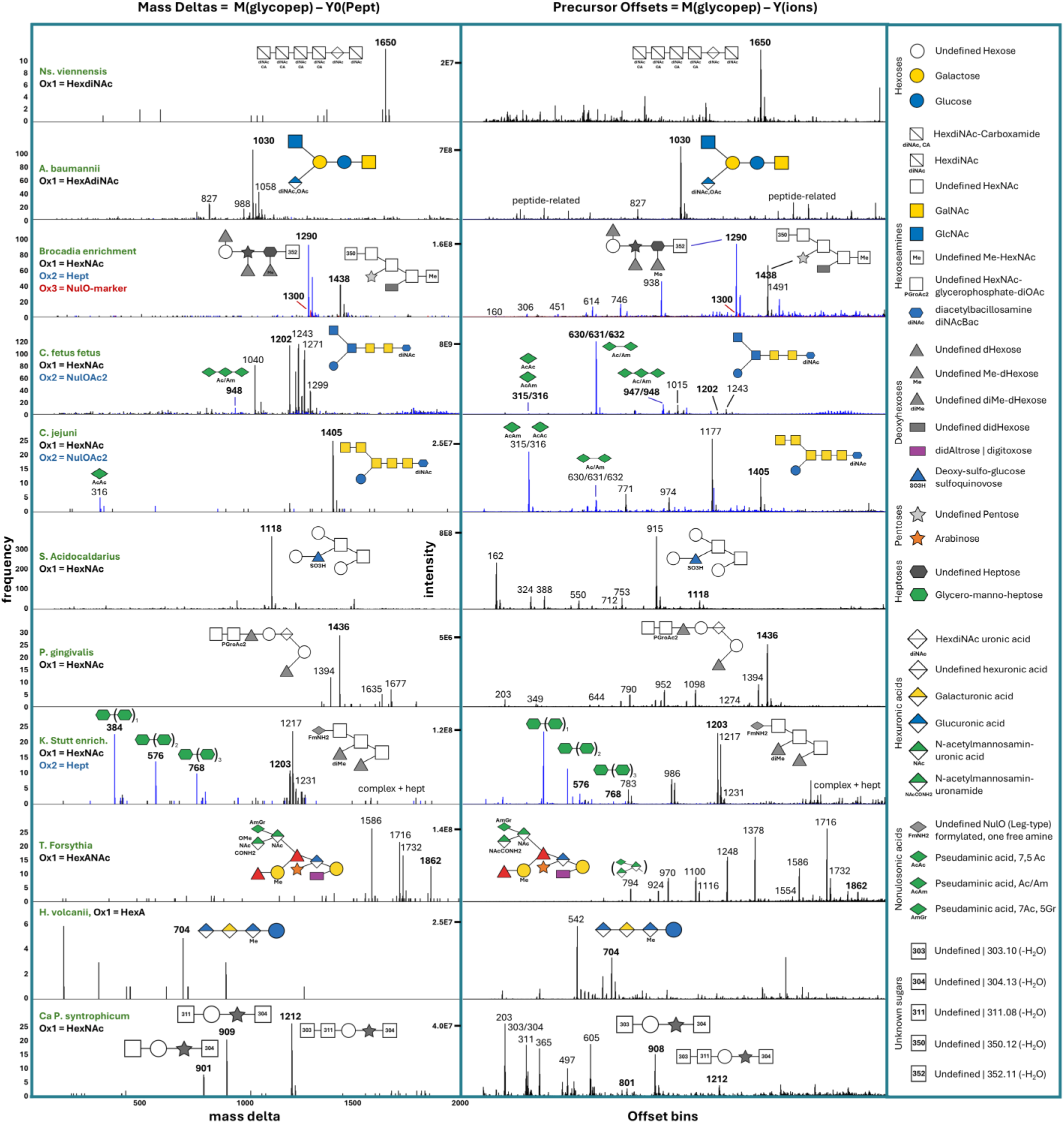
NovoGlycoX glycan profiles across microbial species as obtained from shotgun proteomics data. The histograms display identified glycans derived from spectra containing diagnostic oxonium ions, using two approaches: sequence tag matching (mass delta) and parent mass offsets (offset bins). When multiple oxonium ion clusters are present, one trace per cluster is overlaid in different colours. Both methods yield similar glycan profiles. However, the parent mass offset approach additionally captures fragment ions originating from the reducing end, enabling determination of glycan sequence, linking sugar, and linkage type (N- or O-linked). In contrast, sequence tag matching depends on strong peptide backbone fragmentation to infer glycan masses. For the major glycoforms, representative glycan structures are shown; monosaccharide symbols are explained in the right panel. Details of the identified major glycoforms, including accurate mass, composition, raw mass spectrometry files, reference databases, and corresponding ProteomeXchange project identifiers, are provided in SI Excel Table 1. From top to bottom, the spectra correspond to: Nitrososphaera viennensis, Acinetobacter baumannii, Ca. Brocadia sapporoensis (enrichment), Campylobacter fetus subsp. fetus, Campylobacter jejuni, Sulfolobus acidocaldarius, Porphyromonas gingivalis, Ca. Kuenenia stuttgartiensis, Tannerella forsythia, Haloferax volcanii, and Prometheoarchaeum syntrophicum.

Finally, we analyzed two multi-species enrichment cultures. The first was a Ca. Kuenenia stuttgartiensis enrichment culture whose glycoproteome had been analyzed previously^2^. The culture was highly enriched for Kuenenia (>95%) but represented a mixed microbial sample, with a metagenomic reference database containing more than 200K sequences. Despite the metaproteome-scale dataset, NovoGlycoX accurately identified all previously reported glycoforms within minutes on a standard desktop computer. These included complex-type glycans with terminal nonulosonic acid derivatives, as well as heptose-containing oligosaccharides, both linked to the surface layer protein. The second enrichment culture contained approximately 60% Ca. Brocadia sapporoensis, along with multiple other community members. This planctomycete is phylogenetically related to Ca. K. stuttgartiensis and exhibits the same Me-HexNAc core glycan (1438 m/z) but is modified by other derivatives^2, 9^. OxoniumBrowser identified additional oxonium clusters in this enrichment, for example glycans of 1290 Da (associated with a NulO marker oxonium ion) and 1300 Da (associated with a heptose oxonium ion), which were assigned to co-existing Ignavibacteria strains^2^. Interestingly, the E. coli negative control showed unexpected hexose oxonium ions (see earlier section). As sporadic reports about protein glycosylation in E. coli persist^66-69^, we further investigated the nature of these peaks. Using OxoniumBrowser, we detected low-frequency hexose and dihexose fragment peaks. However, the spectra lacked characteristic sequential 162 Da losses, and the putative hexose-modified peptides eluted late in the gradient, both of which are inconsistent with true glycopeptides. We observed similar patterns in other unrelated microbial strains processed with the same preparation protocol. After further investigating the employed reagents, we found that the bacterial protein extraction reagent contained a glycoside-type detergent. After replacing this reagent, the apparent hexose modifications disappeared. This confirms that the observed oxonium ions were sample preparation adducts, not genuine glycopeptides (SI Doc Figure 4). Overall, this highlights that microbial glycoproteomics is prone to both false negative and false positives if data are not explored carefully.

## CONCLUSION

In summary, we developed NovoGlyco, an open-source platform for microbial glycoproteomics, freely available under the Apache License 2.0. The Python code and Docker versions of both modules, OxoniumBrowser and NovoGlycoX, including dependencies and documentation, are available via SourceForge: https://sourceforge.net/u/glycolab/profile/. The glycoproteomics raw data are summarized in SI Excel Table 1 and are publicly accessible through the ProteomeXchange repositories and the 4TU.ResearchData platform (project “NovoGlyco”): https://data.4tu.nl. Both modules also feature an interactive, browser-based dashboard that allows intuitive exploration of oxonium ions, glycan structures, and glycoproteins.

We demonstrate that OxoniumBrowser enables de novo identification of oxonium ions using either a curated sugar database or an expanded chemical composition space comprising over 3000 plausible compositions. NovoGlycoX further performs oxonium ion guided identification of the glycans and provides detailed insights into glycan sequence, linkage type (O- and N-glycans) and associated glycoproteins. Notably, it uncovered previously unreported flagellin sialylation in Campylobacter fetus, a modification linked to virulence in related Campylobacter jejuni pathogens. The approach is broadly applicable to organisms with multiple glycosylation sites and mixed microbial systems, thereby enabling the exploration of glycosylation in non-model organisms across environmental and clinical contexts.

## Supporting information

SI DOC

## ACKNOWLEDGEMENTS

The authors thank colleagues from the microbial proteomics group for valuable discussions, Dita Heikens for support in the mass spectrometry facility, and Jitske van Ede for providing C. jejuni biomass. The authors further acknowledge all who made their mass spectrometric raw data available through public repositories. D.S. was supported by the MOBODL-2023 grant from HRZZ. Generative AI tools were used for language editing and code optimization. The authors reviewed the content and take full responsibility for the published article.

