## Supplementary material for "NovoGlyco: mapping protein glycosylation in prokaryotes": SI DOC

**Table of contents**

1. OxoniumBrowser architecture, input parameters and output interface
2. NovoGlycoX architecture, input parameters and output interface
3. C. jejuni N-glycan fragmentation profiles at different collision energies (CEs)
4. Lysis buffer induced hexose glycan artefacts in non-glycosylated E. coli
5. C. jejuni and C. fetus NulO flagellin O-glycan fragmentation spectra

#### 1. Oxonium Browser architecture, input parameters and output interface

**Oxonium Browser Architecture.** Oxonium Browser is implemented as a modular Python pipeline (Python 3.11) containerised in Docker for reproducible deployment. As described in the main text, MS/MS spectra are first subjected to a SAGE database search to identify and exclude unmodified peptide spectra. The SAGE database search performs peptide-spectrum matching against the user-provided FASTA database using a target-decoy approach with tryptic digestion (KR/P), carbamidomethylation (C) as a static modification, and oxidation (M) as a variable modification. The default FDR threshold is set at 1%, though this can be adjusted along with other SAGE parameters including minimum peptide length (default: 6), missed cleavages (default: 2), and precursor and fragment mass tolerances (default: 20 ppm each). Scan numbers of identified peptide spectra are collected for exclusion from downstream analysis. Second, a two-pass mass recalibration is applied to all MS2 spectra. In the first pass, seven amino acid fragment reference peaks (147.113, 175.119, 201.123, 215.139, 228.134, 258.145, 292.129 m/z) are matched across all spectra at 20 ppm tolerance, requiring a minimum of 500 matched spectra per reference peak. A global linear calibration function is fitted by least squares regression and applied to all spectra. In the second pass, each spectrum is individually recalibrated at 10 ppm tolerance, requiring at least three matched reference peaks for a stable linear fit. Spectra with insufficient matches retain the global calibration. The resulting mass error distribution plot showing histograms for each of the amino acid reference peaks allows the user to assess calibration quality and select an appropriate mass error tolerance for oxonium ion detection. Third, oxonium ion detection is performed on all spectra not identified as peptides by the SAGE search. As described in the main text, the detection is guided by a curated database of over 35 monosaccharides compiled from CSDB, KEGG, and published glycan literature. The database is user-extensible, and custom monosaccharides can be added. A chemical space database of 3332 theoretical monosaccharide compositions can optionally be enabled, constructed by generating chemically plausible sugar compositions within a defined elemental space (C, H, O, N, S) and mass range (Pabst et al., 2022. The ISME Journal). Each database entry defines two diagnostic fragment masses: the intact oxonium ion ($[M+H-H_2O]^+$) and its water loss fragment ($[M+H-2H_2O]^+$). An equal number of random test masses are included as built-in negative controls, generated within the same mass range (100-400 Da) and cross-checked against real masses, enabling empirical false discovery assessment. For each spectrum, the scanner checks whether both diagnostic masses are present within the defined mass error tolerance. If both masses are found and their average normalized intensity exceeds a user-defined threshold, the detection is recorded. Detection metrics computed per oxonium ion include normalized presence (percentage of spectra containing the ion pair), spectral count, and normalized and raw intensity averages across positive detections. Fourth, results are organized and visualized through an interactive Plotly Dash dashboard. A sortable match table lists all detected oxonium ions grouped by ±18 Da water loss families, with color-coded detection metrics (spectral count, normalized intensity, and spectral presence). A co-occurrence heatmap displays pairwise scan overlap fractions sorted by Jaccard similarity-based hierarchical clustering and accompanied by a dendrogram showing the grouping structure. Retention time profiles with extracted ion chromatograms can be generated for selected ions across the chromatographic run. A comprehensive user guide including pipeline and dashboard documentation, parameter optimization strategies, and detection metrics is available at https://sourceforge.net/p/oxoniumbrowserx/wiki/.
**Configurable Parameters.** The Oxonium Browser offers several key parameters that can be adjusted at runtime to optimize detection performance. The mass error tolerance (MASS_ERROR) defines the maximum allowed mass difference in Daltons between the theoretical and observed oxonium ion mass. The default of 0.001 Da is appropriate for most high-resolution Orbitrap instruments. This can be tightened to 0.0005 Da for increased specificity. For lower-resolution instruments, a larger max mass difference should be set. After processing, the recalibration diagnostic plot reports the average peptide fragment mass error for 90% of reference masses, which can guide the selection of an appropriate tolerance. The intensity threshold (INTENSITY_THRESHOLD, default: 0.25%) sets the minimum normalized intensity for a positive detection, expressed as the average intensity of the diagnostic mass pair relative to total spectrum intensity. For detecting low-abundance or poorly ionising sugars, particularly in bacterial samples, this can be lowered to

0.1-0.15% (or even lower), with test mass performance monitored to assess the impact on false discoveries. For complex samples with dense spectral background, increasing to 0.5-1.0% may reduce random annotations (but also sensitivity). An amino acid marker option (AMINO_ACID_MARKER) can be enabled to require the co-detection of at least one amino acid immonium ion (147.113 or 175.119 m/z, corresponding to lysine and arginine fragments) alongside the oxonium ion pair. This provides an additional validation step, confirming glycopeptide origin. The chemical space search option (CHEMSPACE_SEARCH, default: false) extends the analysis beyond the curated database by additionally scanning against over 3300 systematically enumerated monosaccharide compositions with their own proportional set of random test masses. Both databases are scanned in a single pass, and the interactive dashboard allows switching between curated, chemical space, and combined result views. When viewing chemical space results, stricter default thresholds are automatically applied (counts ≥ 25, intensity ≥ 1.0%, presence ≥ 0.025%) to account for the substantially larger search space. **Output interface.** Supplementary Figure 1 shows a representative Oxonium Browser dashboard output for a yeast whole-cell lysate dataset (S. cerevisiae). At the top, three adjustable threshold filters (spectral count, normalized intensity, spectral presence) control which oxonium ions are displayed across all visualizations. Below the filters, a sortable match table lists detected oxonium ions grouped by ±18 Da water loss families, with color-coded detection metrics. In this example, three sugars pass the default thresholds: Hex (2046 scans, 5.36% intensity, 4.68% presence), HexNAc (544 scans, 7.29% intensity, 1.24% presence), and HexAdiNAc (212 scans, 0.90% intensity, 0.48% presence), with no random test masses detected. Hex and HexNAc are expected components of yeast N-glycans. HexAdiNAc (m/z 259.0925), however, is a nitrogen-rich sugar whose mass overlaps substantially with peptide-derived fragment ions even at very high mass resolution, making it unsuitable as standalone glycan marker. The co-occurrence heatmap and elution profile plot provide additional evidence for evaluating such ambiguous hits. Below the table, the co-occurrence heatmap displays pairwise scan overlap fractions between selected ions, with rows and columns reordered by hierarchical clustering and an accompanying dendrogram. HexNAc and Hex cluster together with high co-occurrence, consistent with their known co-modification on yeast N-glycans, while HexAdiNAc clusters separately with low co-occurrence, supporting a different, likely non-glycan origin. The elution profile plot displays retention time distribution for each selected oxonium ion across the chromatographic run, with Hex and HexNAc showing broad elution consistent with glycopeptide signals. At the bottom, the mass error distribution plot shows that the average peptide fragment mass error for 90% of masses is ± 0.0010 Da, confirming successful recalibration and supporting the use of the default 0.001 Da mass error tolerance for oxonium ion detection.

Dash

127.0.0.1:8051

### Oxonium Browser X

#### Oxonium Browser results for MP_14052020_Yeast_AE16_120min_gps_DDA01

Check Oxonium Browser user guide

counts (default 10-25) > 25 spectral intensity (default 0.1-0.25)> 1.25 spectral presence (default 0.01-0.025) > 0.025

**sugar database:** ○ curated ○ chemspace ◉ both

| group | Oxonium | mass (Da) | count | intensity % | presence % |
|---|---|---|---|---|---|
| 1 | 138.091 | 138.0913 | 573 | 1.3883 | 1.3094 |
| 2 | 144.066 | 144.0655 | 205 | 1.4374 | 0.4685 |
| 3 | 145.050 | 145.0495 | **1560** | 2.2757 | **3.565** |
| 3 | Hex | 163.0601 | **1373** | 2.78 | **3.1376** |
| 3 | 163.060 | 163.0601 | **1374** | 2.7818 | **3.1399** |
| 4 | 168.066 | 168.0655 | 27 | 1.5738 | 0.0617 |
| 4 | 186.076 | 186.0761 | 261 | 2.5763 | 0.5964 |
| 4 | HexNAc | 204.0866 | 260 | **6.4679** | 0.5942 |
| 4 | 204.087 | 204.0867 | 260 | **6.4679** | 0.5942 |
| 4 | 222.097 | 222.0972 | 50 | **6.8226** | 0.1143 |
| 5 | Ox_test_3311 | 322.1879 | 63 | 1.2564 | 0.144 |

export table

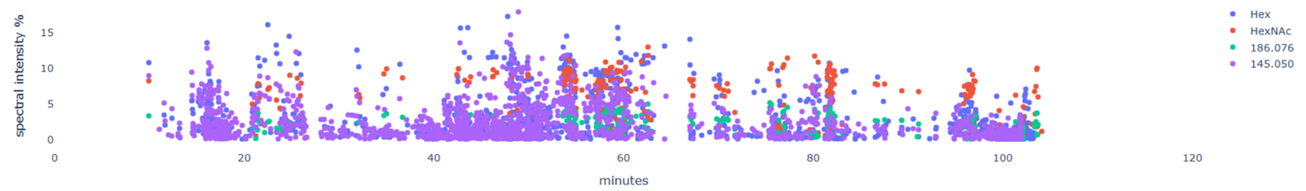


**selected oxonium mass traces**

☑ 145.050 ☐ 163.060 ☑ Hex ☐ 138.091 ☑ 186.076 ☑ HexNAc ☐ 204.087 ☐ 144.066 ☐ 222.097 ☐ 168.066 ☐ Ox_test_3311

export selection

Oxonium Ion Co-occurrence (clustered)

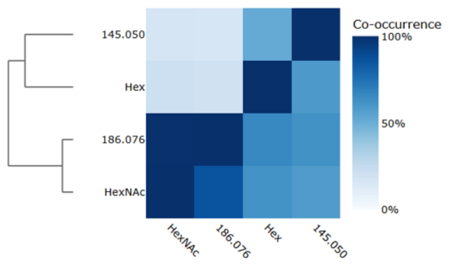


average peptide fragment mass error for 90% of masses: ± 0.0010

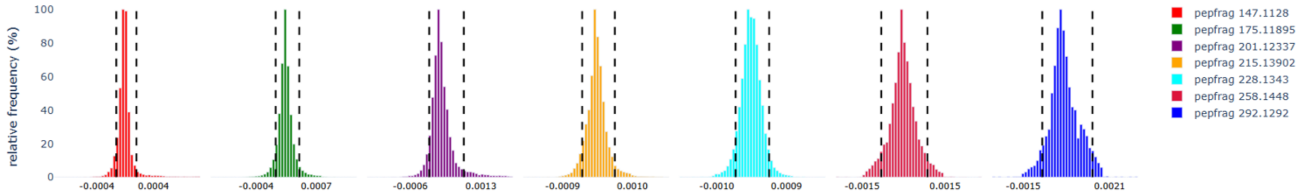

**SI Figure 1. Oxonium Browser dashboard output for a yeast whole-cell lysate dataset (*S. cerevisiae*).** From top to bottom: adjustable threshold filters for spectral count, normalized intensity, and spectral presence; a sortable match table listing detected oxonium ions grouped by ±18 Da water loss families with color-coded metrics showing three entries pass default thresholds (Hex, HexNAc, HexAdiNAc) with no random test masses detected; clustered co-occurrence heatmap showing HexNAc and Hex cluster together consistent with their co-occurrence on yeast N-glycans, while HexAdiNAc clusters separately; elution profiles showing Hex and HexNAc detected throughout the gradient consistent with glycopeptide elution; and mass error distribution histograms for the seven amino acid fragment reference peaks used during two-pass recalibration, reporting an average mass error of ±0.0010 Da for 90% of masses.

#### 2. NovoGlyco architechture, input parameters and ouptut interface

**NovoGlyco Architecture.** NovoGlyco is implemented as a modular Python pipeline (Python 3.11) containerised in Docker for reproducible deployment. As described in the main text, two complementary strategies are applied: sequence tag matching for glycopeptide identification and glycan mass (mass delta) determination, and database-independent precursor mass offset analysis for glycan composition reconstruction. Using the same SAGE configuration as in OxoniumBrowser, MS/MS spectra are first subjected to a conventional database search to identify and exclude unmodified peptide spectra. Identified proteins form a focused database for downstream tag matching, and scan numbers of identified peptide spectra are collected and excluded from the glycopeptide analysis. All remaining unmatched MS2 spectra then undergo charge state deconvolution, where multiply charged fragment ions are converted to singly charged equivalents by detecting isotope spacing patterns for charge states 2-4. After that, the presence of diagnostic oxonium ions from the user-provided list (typically generated by OxoniumBrowser) is annotated for each spectrum. As described in the main text, each oxonium ion requires the co-detection of both the intact ion and its water loss fragment. Intensity values are normalised to total ion current, and spectra containing both diagnostic masses above a user-defined intensity threshold (default: 0.1% of TIC) are flagged. De novo sequence tags are generated from all unmatched spectra using DirecTag, producing up to ten tags per spectrum of user-specified length (default: 5 amino acids), ranked by composite scoring metrics (JointpValue, MzFidelity, Complement, Intensity). Tags are matched against in-silico-generated peptides from the focused protein database, with the option to restrict matches to peptides containing potential glycosylation sites (default: serine, threonine, or asparagine). Valid matches require complete tag presence within the candidate peptide and the peptide mass to be at least 100 Da smaller than the precursor mass. When enabled, $Y_0$ ion validation confirms the presence of the unmodified peptide ion within 20 ppm, with monoisotopic assignment verified by the absence of a peak at -1.0033 Da; for peptides exceeding 2000 Da, doubly charged second or third isotopic peaks are additionally accepted. For each spectrum, the peptide-protein combination supported by the highest number of independently matched tags is reported, with tag count redundancy serving as the primary confidence metric. Precursor mass offsets are then calculated for all processed spectra as the mass difference between the precursor and each decharged fragment ion above a defined mass threshold (default: 750 Da), corresponding to sequential monosaccharide losses from the intact glycopeptide and providing glycan composition information independently of successful peptide identification. Complementary peptide offsets are calculated as mass differences between fragment ions and the $Y_0$ ion, representing sequential monosaccharide additions to the peptide backbone and identifying the linking sugar. Identified glycopeptides are cross-referenced against SAGE results to flag cases where the unmodified peptide counterpart was also detected. Results are organised and visualised through an interactive Plotly Dash dashboard. An oxonium ion co-occurrence heatmap displays pairwise scan overlap fractions sorted by Jaccard similarity-based hierarchical clustering and accompanied by a dendrogram revealing which monosaccharides co-occur in the same spectra. Mass delta and precursor offset histograms with selectable oxonium ion overlays allow comparison of glycan mass distributions across sugar types. Clicking a histogram bin triggers child plots showing complementary offset distributions and m/z spectra, with a second click further narrowing the selection. A glycoprotein candidate table ranks identifications by unique peptide count, with collapsible rows revealing individual peptide-spectrum matches, scoring metrics, and per-scan oxonium annotations. Oxonium filter checkboxes enable progressive refinement from all untargeted results to candidates confirmed by specific sugar fragments. Results can be exported as Excel files and in PeptideShaker-compatible format. A comprehensive user guide including pipeline and dashboard documentation, parameter optimization strategies, and detection metrics is available at https://sourceforge.net/p/novoglycox/wiki/. **Configurable Parameters.** NovoGlyco offers several parameters that can be adjusted at runtime to optimise identification performance. Default values are optimised for high-resolution Orbitrap data from prokaryotic samples. The most critical parameters for glycopeptide identification are the sequence tag length, oxonium ion intensity threshold, and $Y_0$ validation. The sequence tag length (TAG_LENGTH, default: 5) defines the number of amino acids per de novo tag generated by DirecTag. Longer tags require good backbone fragmentation quality but provide more specific database matches, while shorter tags increase sensitivity for spectra with poor peptide fragmentation. The oxonium ion intensity threshold (INT, default: 0.1% of TIC) sets the minimum normalised intensity for a positive oxonium detection, expressed as the average intensity of the diagnostic mass pair relative to total ion current. Values of 0.01-0.05 increase sensitivity for low-abundance glycopeptides, while values of 0.5-1.0 reduce false positives in noisy spectra or samples containing free oligosaccharides. $Y_0$ ion presence validation (VALIDATE_PEPTIDE_MASS, default: true) can be enabled or disabled to control the stringency of glycopeptide identification; disabling it increases sensitivity for samples where $Y_0$ ions are weak or absent, at the cost of potential false positives. The mass error tolerance (MASS_ERROR, default: 0.005 Da) defines the maximum allowed mass difference for oxonium ion pair detection and should be matched to the instrument's mass accuracy, with 0.001 Da appropriate for ultra-high-resolution instruments and 0.01 Da for lower-resolution data. The minimum offset threshold (MIN_OFFSET, default: 750 Da) controls the lower bound for precursor offset calculation and can be lowered to 500-600 Da for small glycans or raised to 900-1000 Da to focus on larger glycan structures. The activation type exclusion (TYPE, default: ETD) removes spectra of the specified fragmentation method from analysis, as HCD typically provides better glycan fragmentation and oxonium ion production. The glycosylation site filter (GLYCOSITES, default: [STN]) restricts candidate peptides to those containing potential glycosylation sites and can be narrowed for specific glycosylation consensus sequons. An amino acid marker option (AMINO_ACID_MARKER, default: false) can be enabled to require co-detection of amino acid immonium ions alongside the oxonium ion pair, confirming glycopeptide origin in samples containing abundant free oligosaccharides. The SAGE search parameters (precursor tolerance, fragment tolerance, missed cleavages, FDR threshold) and visualisation parameters (bin width, port) can also be adjusted. **Output interface.** Supplementary Figure 2 shows a representative NovoGlyco dashboard output for *Acinetobacter baumannii* ATCC 17978, which carries a known pentasaccharide composed of a bacterial-specific di-N-acetylated hexuronic acid (GlcNAc3NAcA4OAc, 300 Da), two HexNAc and two Hex residues. The top panel displays the oxonium ion co-occurrence heatmap, where hierarchical clustering reveals that HexAdiNAc, NulONH2, and NeuAc (fragmentation products of the GlcNAc3NAcA4OAc) frequently co-occur with HexNAc, indicating a shared glycan origin. Below the heatmap, the mass delta histogram reveals a dominant peak at 1030.35 Da corresponding to the intact pentasaccharide (GalNAc-Glc-[GlcNAc-]Gal-GlcNAc3NAcA4OAc), along with its methylated and acetylated derivatives at neighbouring masses. Selecting the 1030.30-1030.40 Da bin triggers the child plots shown in the middle panels. The first row shows precursor offsets as paired frequency histogram (left) and summed intensity bar plot (right), revealing HexNAc-Hex-Hex loss series consistent with stepwise removal of the GalNAc linking sugar followed by the two core hexoses. The second row shows peptide offsets in the same paired format, revealing sequential monosaccharide additions to the peptide backbone starting from the $Y_0$ ion. The first peptide offset peak corresponds to a HexNAc addition, confirming N-acetylhexosamine as the linking sugar, again followed by offsets corresponding to two hexoses. The third row shows the binned m/z array distribution with the diagnostic sugar ions, where the most abundant peak corresponds to the GlcNAc3NAcA4OAc oxonium ion at 301 m/z, with its HexAdiNAc fragment at 259 m/z and the HexNAc oxonium ion at 204 m/z also clearly visible. At the bottom, the glycoprotein candidate table lists identified glycoproteins ranked by supporting evidence. Expanding a protein row reveals the individual peptide-spectrum matches with DirecTag scoring metrics and oxonium information.

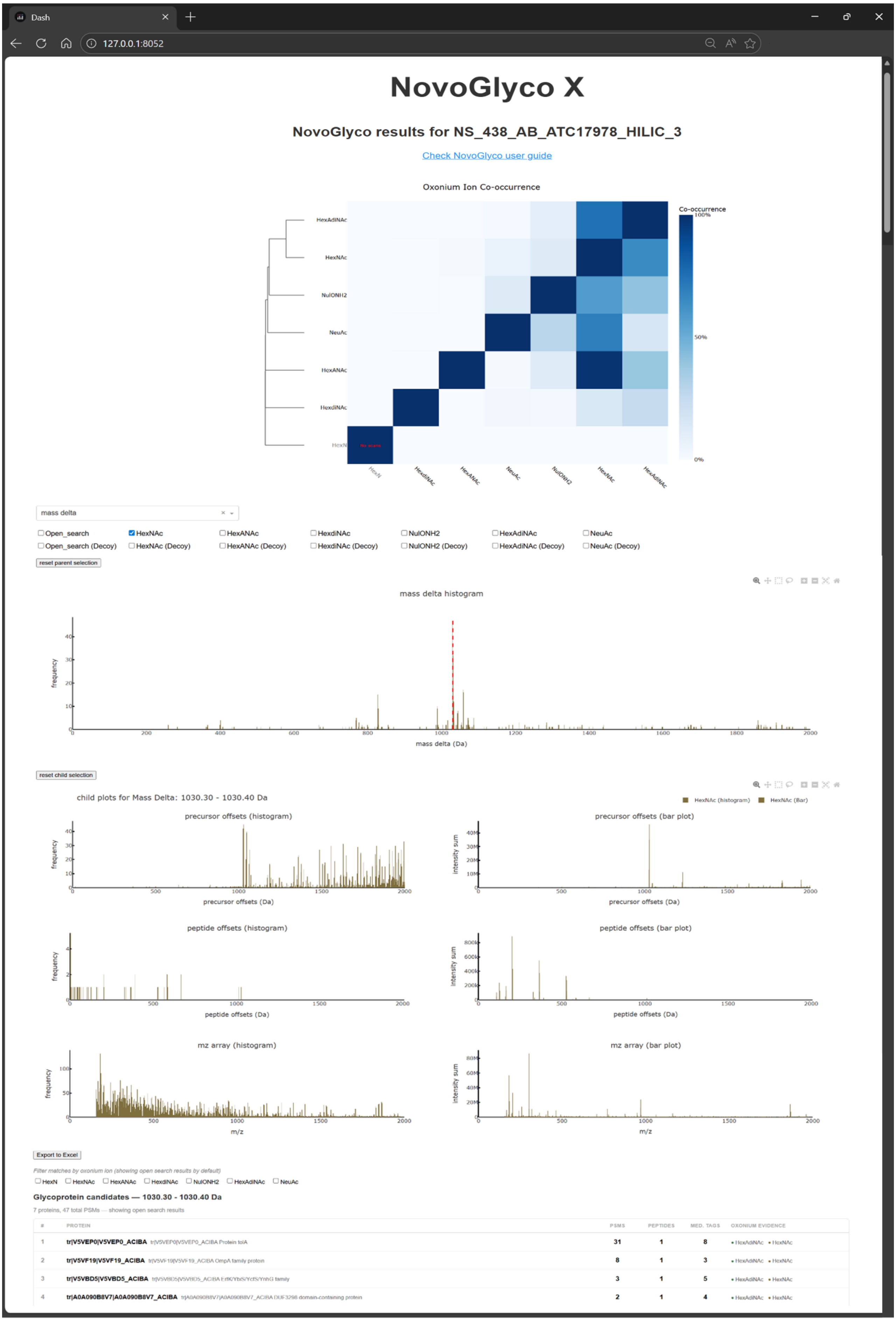
Dash
127.0.0.1:8052
NovoGlyco X
NovoGlyco results for NS_438_AB_ATC17978_HILIC_3
Check NovoGlyco user guide
Oxonium Ion Co-occurrence
Co-occurrence
HexAdiNAc
HexNAc
NulONH2
NeuAc
HexANAc
HexdiNAc
HexN
mass delta
Open_search
HexNAc
HexANAc
HexdiNAc
NulONH2
HexAdiNAc
NeuAc
Open_search (Decoy)
HexNAc (Decoy)
HexANAc (Decoy)
HexdiNAc (Decoy)
NulONH2 (Decoy)
HexAdiNAc (Decoy)
NeuAc (Decoy)
reset parent selection
mass delta histogram
frequency
mass delta (Da)
reset child selection
child plots for Mass Delta: 1030.30 - 1030.40 Da
HexNAc (histogram)
HexNAc (Bar)
precursor offsets (histogram)
precursor offsets (bar plot)
precursor offsets (Da)
intensity sum
peptide offsets (histogram)
peptide offsets (bar plot)
peptide offsets (Da)
mz array (histogram)
mz array (bar plot)
m/z
Export to Excel
Filter matches by oxonium ion (showing open search results by default)
HexN
HexNAc
HexANAc
HexdiNAc
NulONH2
HexAdiNAc
NeuAc
Glycoprotein candidates — 1030.30 - 1030.40 Da
7 proteins, 47 total PSMs — showing open search results
### PROTEIN PSMS PEPTIDES MED. TAGS OXONIUM EVIDENCE
1 tr|V5VEP0|V5VEP0_ACIBA 31 1 8 HexAdiNAc HexNAc
2 tr|V5VF19|V5VF19_ACIBA 8 1 3 HexAdiNAc HexNAc
3 tr|V5VBD5|V5VBD5_ACIBA 3 1 5 HexAdiNAc HexNAc
4 tr|A0A090B8V7|A0A090B8V7_ACIBA 2 1 4 HexAdiNAc HexNAc

**SI Figure 2. NovoGlycoX dashboard output for *Acinetobacter baumannii* ATCC 17978.** From top to bottom: oxonium ion co-occurrence heatmap with Jaccard similarity-based hierarchical clustering and dendrogram, grouping HexAdiNAc, NulONH2, and NeuAc (fragmentation products of GlcNAc3NAcA4OAc) together with HexNAc; mass delta histogram showing the distribution of glycan masses, with a dominant peak at 1030 Da corresponding to the characteristic pentasaccharide and its methylated and acetylated derivatives; child plots triggered by the selected mass delta bin (1030.30-1030.40 Da), shown as paired frequency histograms (left) and summed intensity bar plots (right); glycoprotein candidate table listing proteins ranked by supporting evidence. Precursor offsets reveal a HexNAc-Hex-Hex loss series from the intact glycopeptide, while peptide offsets complementarily show sequential monosaccharide additions starting from the $Y_0$ ion, with HexNAc as the linking sugar followed by two hexoses. The m/z array shows the GlcNAc3NAcA4OAc oxonium ion (301 m/z) as the most abundant peak, with the HexAdiNAc fragment (259 m/z) and HexNAc (204 m/z) also clearly visible. The top-ranked glycoprotein candidate (V5VEP0, Protein NlpA) is supported by 31 PSMs with a median tag count of 8.

#### 3. C. jejuni N-glycan fragmentation profiles at different CEs

**Dependency of mass-tag and precursor offset on collision energy.** The two complementary NovoGlyco strategies described above have inherently different fragmentation requirements: sequence tag matching depends on peptide backbone b/y-ion coverage for successful de novo tag generation, while precursor mass offset analysis relies on the preservation of glycopeptide Y-ion series reflecting sequential monosaccharide losses. To evaluate how collision energy affects each strategy, we analysed glycoproteomics data from *Campylobacter jejuni*, which carries a well-characterised N-linked heptasaccharide (1405.55 Da) composed of diNAcBAc, a hexose, and five N-acetylhexosamines. The same sample was analysed sequentially using discrete normalized collision energies (NCE 28, 33, and 38) and stepped HCD (NCE 20/30/40), which applies multiple collision energies within a single scan cycle to produce hybrid spectra containing both peptide backbone fragments and glycan-derived signals. As shown in Supplementary Figure 3, increasing collision energy had opposing effects on the two strategies. For the first strategy, the mass delta frequency (top row) increased with higher NCE, reflecting improved peptide backbone fragmentation and consequently more sequence tag matches, as the number of identified glycopeptides almost doubled from NCE 28 to NCE 38. This improvement, however, came at the direct cost of glycan structural information in the second strategy. The precursor offset patterns (bottom row) show that at NCE 28, the complete sequential monosaccharide series is observed (from the linking diNAcBAc through successive HexNAc and terminal Hex losses), enabling full reconstruction of the glycan sequence. At NCE 33, the lower offset peaks begin to diminish as intermediate glycopeptide Y-ions are lost to secondary fragmentation. At NCE 38, elevated fragmentation energy led to near-complete dissociation of both peptide and glycan moieties, retaining only the highest-mass offsets and substantially reducing the information content of the precursor offset analysis. This illustrates a fundamental tension between the two strategies: conditions that maximise peptide identification simultaneously diminish glycan composition analysis. Stepped HCD resolved this trade-off by providing hybrid spectra in which both strategies operate effectively, providing reliable peptide identification while preserving sufficient glycan structural information. Mass delta frequencies were comparable to NCE 33, indicating robust peptide identification, while the full precursor offset ladder remained intact with nearly all sequential monosaccharide losses clearly resolved. Thus, collision energy selection directly impacts the complementarity of the two NovoGlyco analytical strategies, with stepped HCD generally being recommended for comprehensive glycopeptide characterisation where both peptide identity and glycan composition are needed.

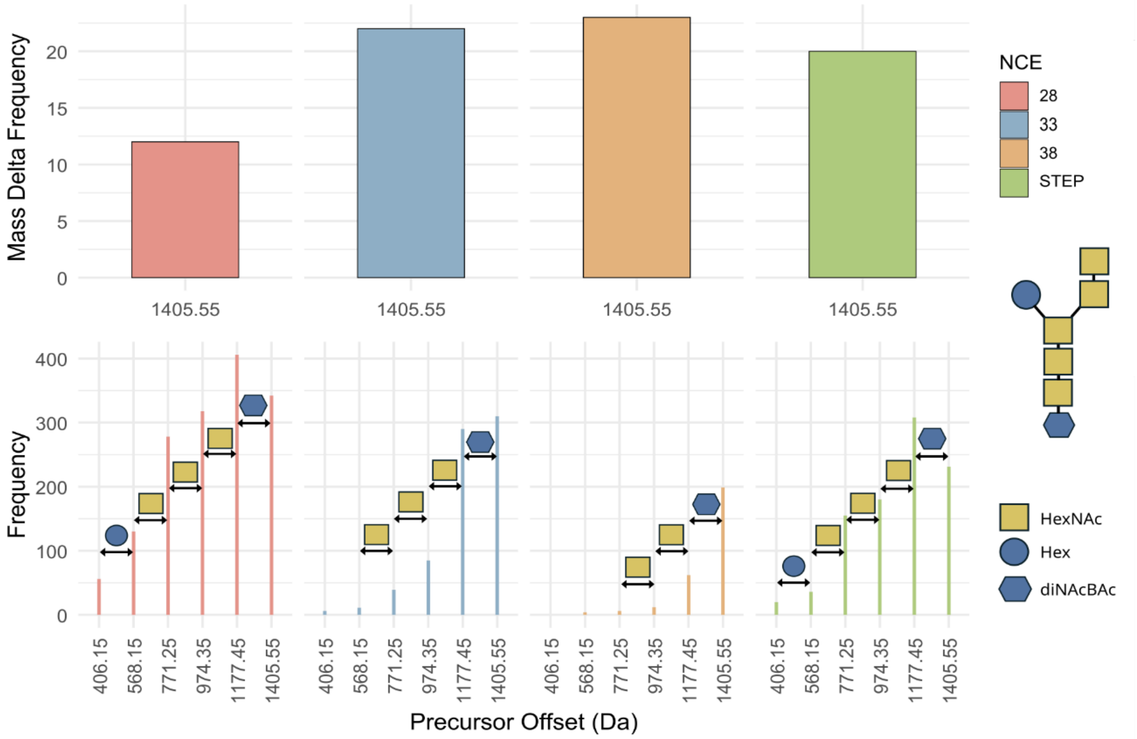


**SI Figure 3. Effect of normalized collision energy (NCE) on NovoGlyco analysis of *C. jejuni* N-glycopeptides carrying the well-studied heptasaccharide (1405.55 Da).** Mass delta distributions (top row) and precursor offset patterns (bottom row) are compared across different fixed collision energies (NCE 28, 33, 38) and stepped HCD (NCE 20/30/40). Monosaccharide assignments are annotated on the offset peaks: Hex (circle), HexNAc (square), and diNAcBAc (hexagon). Higher collision energies increase mass delta frequency (more successful peptide identifications), but progressively diminish the precursor offset ladder used for glycan characterization. Stepped HCD preserves the complete glycan fragmentation series while maintaining high peptide identification rates.

#### 4. Lysis buffer induced hexose glycan artefact demonstrated with non-glycosylated E. Coli K12

**a)** **Oxonium Browser results for MP_DS_04072024_bper_acetone_120mins_gps_DDA01**

Check Oxonium Browser user guide

counts (default 10-25) > 5 spectral intensity (default 0.1-0.25)> 1 spectral presence (default 0.01-0.025) > 0

| group | Oxonium | mass (Da) | count | intensity % | presence % |
|---|---|---|---|---|---|
| 1 | Hex | 163.0601 | 7 | 4.3512 | 0.0173 |

**b)** **Oxonium Browser results for MP_DS_04072024_abc_acetone_120mins_gps_DDA01**

Check Oxonium Browser user guide

counts (default 10-25) > 5 spectral intensity (default 0.1-0.25)> 1 spectral presence (default 0.01-0.025) > 0

**SI Figure 4. B-PER–derived hexose fragment artifacts in non-glycosylated E. coli K12.** (a) Samples lysed with B-PER buffer exhibit a series of hexose oxonium ions that co-elute and co-fragment with peptides, as identified using OxoniumBrowser (oxonium intensity threshold = 0.1% rel. abundance; amino acid marker validation enabled). With amino acid validation active, hexose-derived oxonium ions are co-isolated and fragmented alongside peptide ions. (b) In contrast, samples prepared using ammonium bicarbonate and sodium deoxycholate buffers show no detectable hexose fragments. These B-PER–derived artifacts can be misinterpreted as glycopeptide spectra but are fully eliminated upon removal of B-PER from the workflow, consistent with the presence of a hexose-based alkyl glucoside surfactant.

#### 5. Fragmentation spectra novel glycans

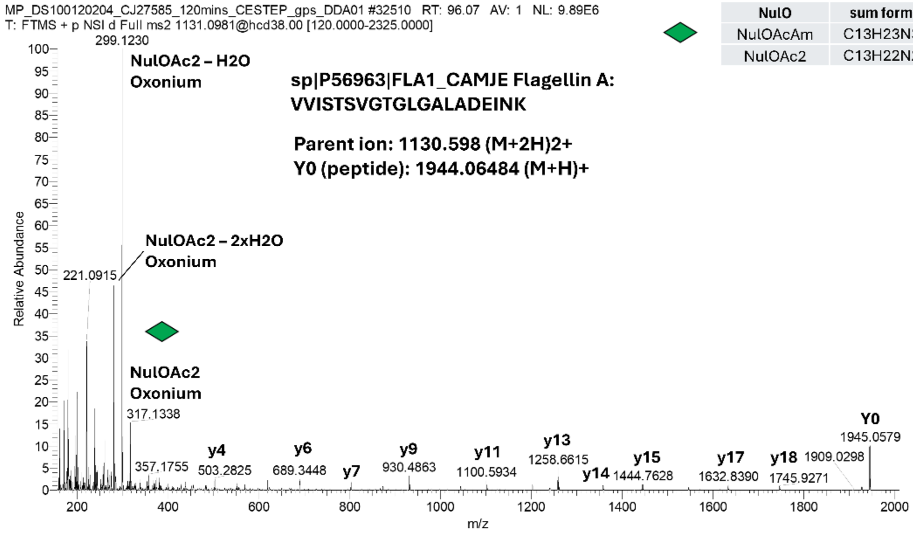


| NulO | sum formula | mi | mi - $H_2O$ | mi - $H_2O$ + H+ |
|---|---|---|---|---|
| NulOAcAm | C13H23N3O7 | 333.1536 | 315.14304 | 316.15024 |
| NulOAc2 | C13H22N2O8 | 334.13762 | 316.12706 | 317.13426 |

**Fragment Ion Table, monoisotopic masses**

| Seq | # | B | Y | # (+1) |
|---|---|---|---|---|
| V | 1 | 100.07574 | 1944.06484 | 20 |
| V | 2 | 199.14415 | 1844.99642 | 19 |
| I | 3 | 312.22821 | 1745.92801 | 18 |
| S | 4 | 399.26024 | 1632.84395 | 17 |
| T | 5 | 500.30792 | 1545.81192 | 16 |
| S | 6 | 587.33995 | 1444.76424 | 15 |
| V | 7 | 686.40836 | 1357.73221 | 14 |
| G | 8 | 743.42982 | 1258.66380 | 13 |
| T | 9 | 844.47750 | 1201.64233 | 12 |
| G | 10 | 901.49897 | 1100.59466 | 11 |
| L | 11 | 1014.58303 | 1043.57319 | 10 |
| G | 12 | 1071.60449 | 930.48913 | 9 |
| A | 13 | 1142.64161 | 873.46767 | 8 |
| L | 14 | 1255.72567 | 802.43055 | 7 |
| A | 15 | 1326.76278 | 689.34649 | 6 |
| D | 16 | 1441.78973 | 618.30938 | 5 |
| E | 17 | 1570.83232 | 503.28243 | 4 |
| I | 18 | 1683.91638 | 374.23984 | 3 |
| N | 19 | 1797.95931 | 261.15578 | 2 |
| K | 20 | 1926.05427 | 147.11285 | 1 |

**SI Figure 5. C. jejuni FlaA O-glycosylation.** The fragmentation spectrum shows the C. jejuni NuloAcAc-type O-glycans (single NulO) on the FlaA protein (accession number: sp|P56963|FLA1_CAMJE Flagellin A) with the peptide sequence: VVISTSVGTGLGALADEINK. Fragment ions were calculated using the MS/MS Fragment Ion Calculator: https://db.systemsbiology.net/proteomicsToolkit/FragIonServlet.html.

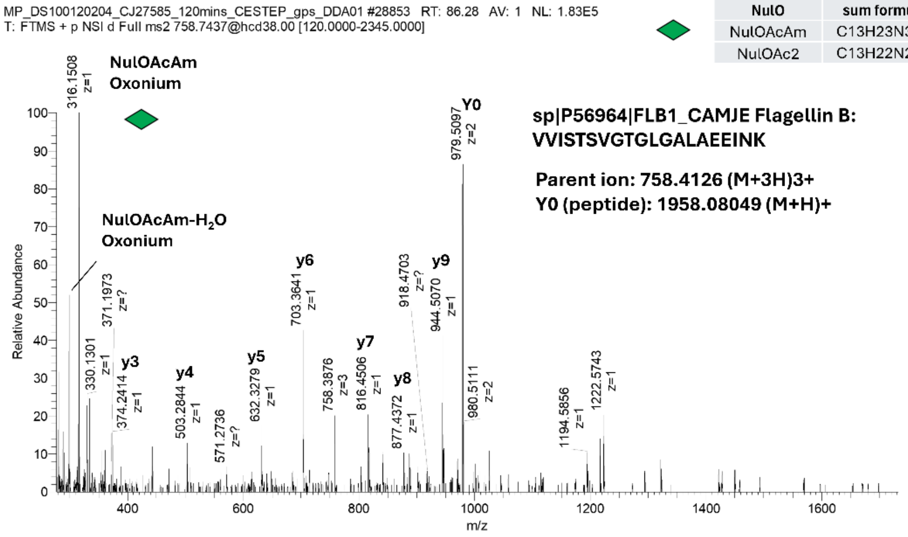


| NulO | sum formula | mi | mi - $H_2O$ | mi - $H_2O$ + H+ |
|---|---|---|---|---|
| NulOAcAm | C13H23N3O7 | 333.1536 | 315.14304 | 316.15024 |
| NulOAc2 | C13H22N2O8 | 334.13762 | 316.12706 | 317.13426 |

**Fragment Ion Table, monoisotopic masses**

| Seq | # | B | Y | # (+1) |
|---|---|---|---|---|
| V | 1 | 100.07574 | 1958.08049 | 20 |
| V | 2 | 199.14415 | 1859.01207 | 19 |
| I | 3 | 312.22821 | 1759.94366 | 18 |
| S | 4 | 399.26024 | 1646.85960 | 17 |
| T | 5 | 500.30792 | 1559.82757 | 16 |
| S | 6 | 587.33995 | 1458.77989 | 15 |
| V | 7 | 686.40836 | 1371.74786 | 14 |
| G | 8 | 743.42982 | 1272.67945 | 13 |
| T | 9 | 844.47750 | 1215.65798 | 12 |
| G | 10 | 901.49897 | 1114.61031 | 11 |
| L | 11 | 1014.58303 | 1057.58884 | 10 |
| G | 12 | 1071.60449 | 944.50478 | 9 |
| A | 13 | 1142.64161 | 887.48332 | 8 |
| L | 14 | 1255.72567 | 816.44620 | 7 |
| A | 15 | 1326.76278 | 703.36214 | 6 |
| E | 16 | 1455.80538 | 632.32503 | 5 |
| E | 17 | 1584.84797 | 503.28243 | 4 |
| I | 18 | 1697.93203 | 374.23984 | 3 |
| N | 19 | 1811.97496 | 261.15578 | 2 |
| K | 20 | 1940.06992 | 147.11285 | 1 |

**SI Figure 6. C. jejuni FlaB O-glycosylation.** The fragmentation spectrum shows the C. jejuni NuloAcAm-type O-glycans (single NulO) on the FlaB protein (accession number: sp|P56964|FLB1_CAMJE Flagellin B) with the peptide sequence: VVISTSVGTGLGALAEEINK. Fragment ions were calculated using the MS/MS Fragment Ion Calculator: https://db.systemsbiology.net/proteomicsToolkit/FragIonServlet.html.

| NulO | sum formula | mi | mi - $H_2O$ | mi - $H_2O$ + H+ |
|---|---|---|---|---|
| NulOAcAm | C13H23N3O7 | 333.1536 | 315.14304 | 316.15024 |
| NulOAc2 | C13H22N2O8 | 334.13762 | 316.12706 | 317.13426 |

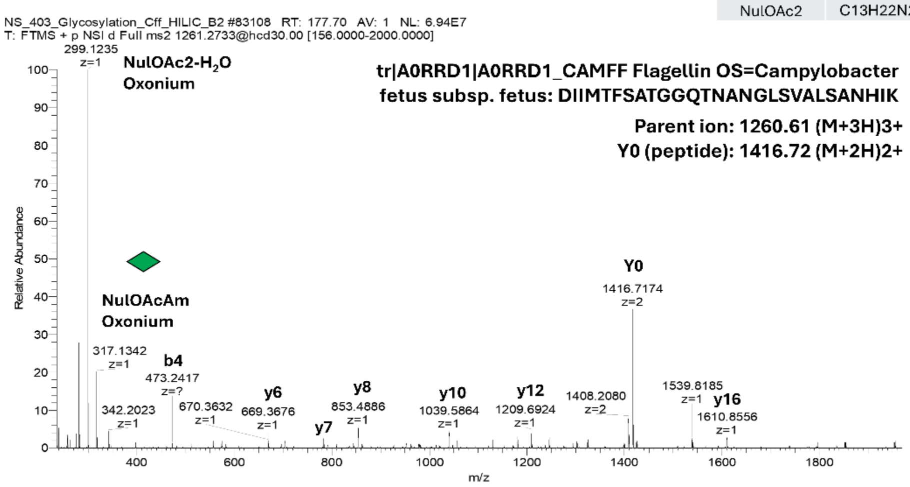


**Fragment Ion Table, monoisotopic masses**

| Seq | # | B | Y | # (+1) |
|---|---|---|---|---|
| D | 1 | 116.03426 | 2831.43566 | 28 |
| I | 2 | 229.11833 | 2716.40871 | 27 |
| I | 3 | 342.20239 | 2603.32465 | 26 |
| M | 4 | 473.24288 | 2490.24059 | 25 |
| T | 5 | 574.29056 | 2359.20010 | 24 |
| F | 6 | 721.35897 | 2258.15242 | 23 |
| S | 7 | 808.39100 | 2111.08401 | 22 |
| A | 8 | 879.42811 | 2024.05198 | 21 |
| T | 9 | 980.47579 | 1953.01487 | 20 |
| G | 10 | 1037.49725 | 1851.96719 | 19 |
| G | 11 | 1094.51872 | 1794.94573 | 18 |
| Q | 12 | 1222.57729 | 1737.92426 | 17 |
| T | 13 | 1323.62497 | 1609.86569 | 16 |
| N | 14 | 1437.66790 | 1508.81801 | 15 |
| A | 15 | 1508.70501 | 1394.77508 | 14 |
| N | 16 | 1622.74794 | 1323.73797 | 13 |
| G | 17 | 1679.76940 | 1209.69504 | 12 |
| L | 18 | 1792.85347 | 1152.67358 | 11 |
| S | 19 | 1879.88549 | 1039.58951 | 10 |
| V | 20 | 1978.95391 | 952.55748 | 9 |
| A | 21 | 2049.99102 | 853.48907 | 8 |
| L | 22 | 2163.07509 | 782.45196 | 7 |
| S | 23 | 2250.10711 | 669.36789 | 6 |
| A | 24 | 2321.14423 | 582.33586 | 5 |
| N | 25 | 2435.18715 | 511.29875 | 4 |
| H | 26 | 2572.24607 | 397.25582 | 3 |
| I | 27 | 2685.33013 | 260.19691 | 2 |
| K | 28 | 2813.42509 | 147.11285 | 1 |

**SI Figure 7. C. fetus FlaA O-glycosylation.** The fragmentation spectrum shows the C. fetus NuloAcAc-type O-glycans (3xNulO) on the Fla protein (accession number: tr|A0RRD1|A0RRD1_CAMFF Flagellin OS=Campylobacter fetus subsp. fetus) with the peptide sequence: DIIMTFSATGGQTNANGLSVALSANHIK. Fragment ions were calculated using the MS/MS Fragment Ion Calculator: https://db.systemsbiology.net/proteomicsToolkit/FragIonServlet.html.

| NulO | sum formula | mi | mi - $H_2O$ | mi - $H_2O$ + H+ |
|---|---|---|---|---|
| NulOAcAm | C13H23N3O7 | 333.1536 | 315.14304 | 316.15024 |
| NulOAc2 | C13H22N2O8 | 334.13762 | 316.12706 | 317.13426 |

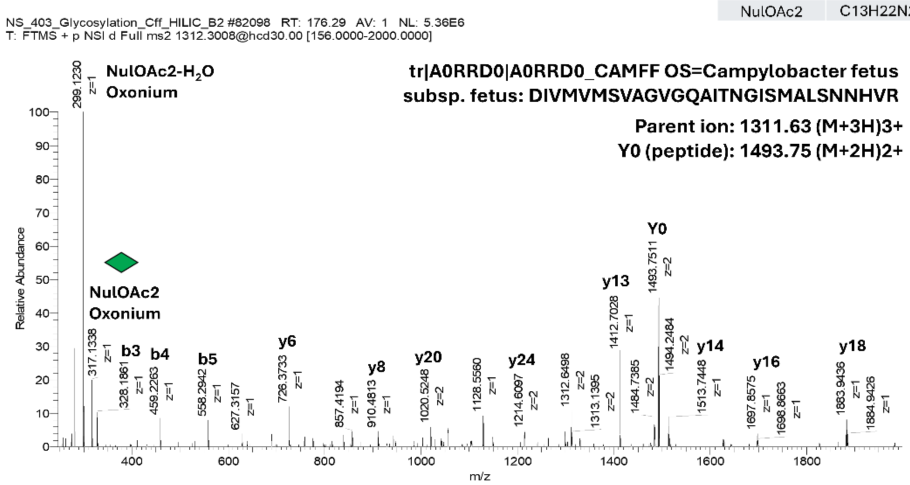


**Fragment Ion Table, monoisotopic masses**

| Seq | # | B | Y | # (+1) |
|---|---|---|---|---|
| D | 1 | 116.03426 | 2984.51148 | 29 |
| I | 2 | 229.11833 | 2869.48454 | 28 |
| V | 3 | 328.18674 | 2756.40048 | 27 |
| M | 4 | 459.22723 | 2657.33206 | 26 |
| V | 5 | 558.29564 | 2526.29158 | 25 |
| M | 6 | 689.33613 | 2427.22316 | 24 |
| S | 7 | 776.36815 | 2296.18268 | 23 |
| V | 8 | 875.43657 | 2209.15065 | 22 |
| A | 9 | 946.47368 | 2110.08224 | 21 |
| G | 10 | 1003.49514 | 2039.04512 | 20 |
| V | 11 | 1102.56356 | 1982.02366 | 19 |
| G | 12 | 1159.58502 | 1882.95525 | 18 |
| Q | 13 | 1287.64360 | 1825.93378 | 17 |
| A | 14 | 1358.68071 | 1697.87521 | 16 |
| I | 15 | 1471.76478 | 1626.83809 | 15 |
| T | 16 | 1572.81245 | 1513.75403 | 14 |
| N | 17 | 1686.85538 | 1412.70635 | 13 |
| G | 18 | 1743.87685 | 1298.66342 | 12 |
| I | 19 | 1856.96091 | 1241.64196 | 11 |
| S | 20 | 1943.99294 | 1128.55790 | 10 |
| M | 21 | 2075.03342 | 1041.52587 | 9 |
| A | 22 | 2146.07054 | 910.48538 | 8 |
| L | 23 | 2259.15460 | 839.44827 | 7 |
| S | 24 | 2346.18663 | 726.36420 | 6 |
| N | 25 | 2460.22956 | 639.33218 | 5 |
| N | 26 | 2574.27248 | 525.28925 | 4 |
| H | 27 | 2711.33139 | 411.24632 | 3 |
| V | 28 | 2810.39981 | 274.18741 | 2 |
| R | 29 | 2966.50092 | 175.11900 | 1 |

**SI Figure 8. C. fetus FlaA O-glycosylation.** The fragmentation spectrum shows the C. fetus NuloAcAc-type O-glycans (3xNulO) on the Fla protein (accession number: tr|A0RRD0|A0RRD0_CAMFF OS=Campylobacter fetus subsp. fetus) with the peptide sequence: DIVMVMSVAGVGQAITNGISMALSNNHVR. Fragment ions were calculated using the MS/MS Fragment Ion Calculator: https://db.systemsbiology.net/proteomicsToolkit/FragIonServlet.html.

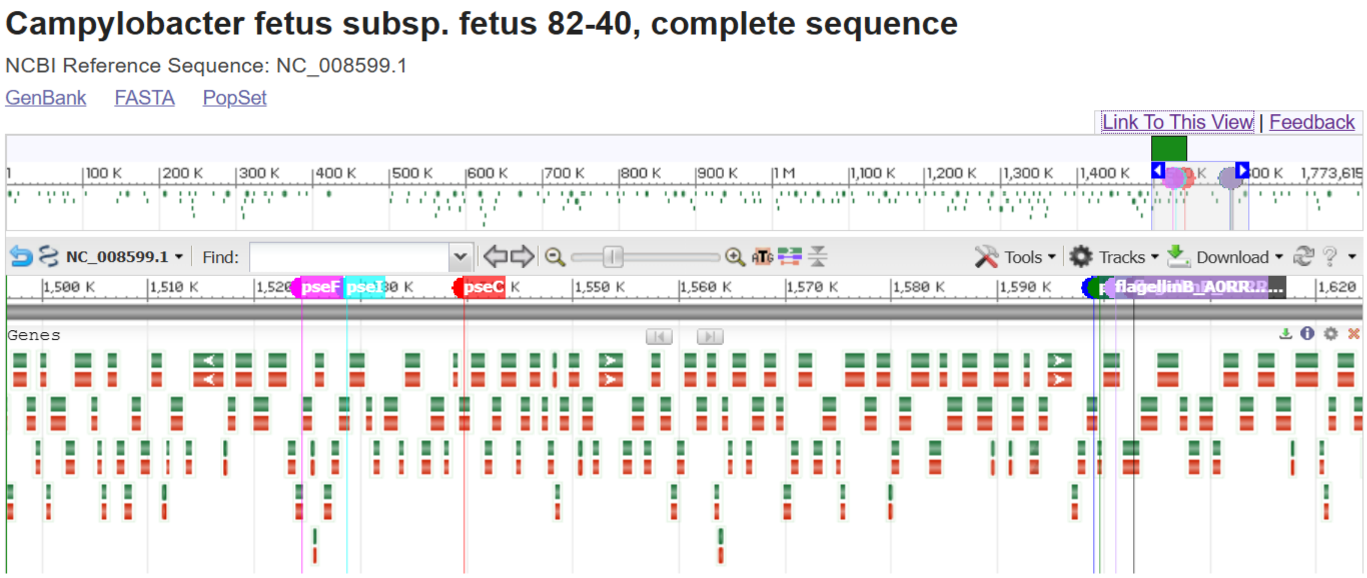


**SI Figure 9. Genomic context of NulO biosynthesis genes in *Campylobacter fetus subsp. fetus* 82-40.** All major pseudaminic acid Pse biosynthesis genes (pseB, pseC, pseH, pseI, and pseF) are readily annotated in the reference genome (NC_008599.1), whereas pseG is not clearly annotated. The identified genes are organized within the same genomic region, with pseB and pseH located in close proximity, and pseI and pseF forming a second nearby locus. Notably, these loci are positioned adjacent to the flagellin genes (CFF8240_RS07985 and CFF8240_RS07990), which were identified here as NulO modified. This genomic organization supports a functional link between Pse biosynthesis and flagellin modification. Legionaminic acid biosynthesis pathway genes were not found to be annotated in the genome NC_008599.1, indicating that the produced NulOs are PseAc2 molecules. The NCBI reference genome NC_008599.1 can be accessed via NCBI: https://www.ncbi.nlm.nih.gov/nuccore/NC_008599.1?report=graph#
